# Genome-wide admixture is common across the *Heliconius* radiation

**DOI:** 10.1101/414201

**Authors:** Krzysztof M. Kozak, W. Owen McMillan, Mathieu Joron, Christopher D. Jiggins

**Affiliations:** Smithsonian Tropical Research Institute, Gamboa, Panama; Department of Zoology, University of Cambridge, Cambridge, United Kingdom of Great Britain and Northern Ireland; Centre d’Ecologie et Evolution, Universite de Montpellier, Montpellier, France

## Abstract

How frequent is gene flow between species? The pattern of evolution is typically portrayed as a phylogenetic tree, implying that speciation is a series of splits between lineages. Yet gene flow between good species is increasingly recognized as an important mechanism in the diversification of radiations, often spreading adaptive traits and leading to a complex pattern of phylogenetic incongruence. This process has thus far been studied in cases involving few species, or geographically restricted to spaces like islands, but not on the scale of a continental radiation. Previous studies have documented gene flow, adaptive introgression and hybrid speciation in a small subsection of the charismatic Neotropical butterflies *Heliconius*. Using genome-wide resequencing of 40 out of 45 species in the genus we demonstrate for the first time that admixture has played a role throughout the evolution of *Heliconius* and the sister genus *Eueides*. Modelling of phylogenetic networks based on 6848 orthologous autosomal genes (Maximum Pseudo-Likelihood Networks) or 5,483,419 high quality SNPs (Ancestral Recombination Graph) uncovers nine new cases of interspecific gene flow at up to half of the genome. However, *f4* statistics of admixture show that the extent of the process has varied between subgenera. Evidence for introgression is found at all five loci controlling the colour and shape of the mimetic wing patterns, including in the putative hybrid species *H. hecalesia*, characterised by an unusual hindwing. Due to hybridization and incomplete coalescence during rapid speciation, individual gene trees show rampant discordance. Although reduced gene flow and faster coalescence are expected at the Z chromosome, we discover high levels of conflict between the 416 sex-linked loci. Despite this discordant pattern, both concatenation and multispecies coalescent approaches yield surprisingly consistent and fully supported genome-wide phylogenies. We conclude that the imposition of the bifurcating tree model without testing for interspecific gene flow may distort our perception of adaptive radiations and thus the ability to study trait evolution in a comparative framework.

## INTRODUCTION

Interspecific hybridization and the resulting gene flow across porous species barriers are increasingly recognized as major processes in evolution, detectable at many taxonomic levels (Abbott et al. 2013; Abbott et al. 2016; Schumer et al. 2014; Feliner et al. 2017). This conceptual shift has been facilitated by whole-genome sequencing yielding precise quantitative estimates of introgression (Schrider et al. 2018). However, there are other factors that create the pattern of phylogenetic discordance in genomic data including retention of ancient polymorphism and population structure, incomplete lineage sorting, selection, gene shuffling and duplication. Phylogenomics has massively expanded the potential of comparative molecular biology to elucidate these phenomena in both ancient clades [e.g. birds (Jarvis et al. 2014); mammals (Song et al. 2012)] and recently diverged taxa [e.g. *Virentes* oaks (Eaton et al. 2015), *Drosophila simulans* (Garrigan et al. 2013), *Danaus plexippus* butterflies (Zhang et al. 2014)], but less so at the intermediate level of large, recently-radiated genera. The few genomic studies of recent radiations published to date confirm the prediction of high discordance among independent markers. Analyses of 16 *Anopheles* species and all 48 families of birds (Jarvis et al. 2014) found that no individual gene tree, whether based on protein-coding sequences or large sliding windows, is identical to the species tree topology. This phenomenon appears to be partially attributable to incomplete lineage sorting (ILS) in rapidly radiating groups exposed to novel ecological opportunities, including Darwin’s finches speciating after the colonisation of the Galapagos archipelago (Lamichhaney et al. 2015), mammals in response to climatic conditions (Song et al. 2012), and birds after the K/T extinction event (Jarvis et al. 2014; Jarvis 2016).

Discordance due to gene flow between species that are not fully reproductively isolated has been detected in several cases, ranging from humans (Meyer et al. 2012; Sankararaman et al. 2016) to sunflowers (Renaut et al. 2014). In many such taxa genome-wide studies demonstrate both stochastic gene flow across the species barrier, as well as adaptive introgression, where natural selection acts to fix introgressed variants. Alleles at a major locus contributing to the variation in beak shape among Darwin’s finches have introgressed between multiple species (Lamichhaney et al. 2015). Similarly, selection has driven a block of transcription factor binding sites from the ecological generalist *Drosophila simulans* to the specialist *D. secchelia* (Brand et al. 2013), and recent malaria prevention efforts have caused a resistance allele to sweep across *Anopheles gambiae* populations and into *A. colluzzi* (Clarkson et al. 2014; Miles et al. 2017). Nonetheless, the extent and importance of hybridisation in fueling speciation remain hotly contested (Schumer et al. 2014; Feliner et al. 2017; Schumer et al. 2018). Furthermore, gene flow has been less frequently considered than ILS in phylogenetic studies of radiations. One reason is perhaps that modelling gene flow poses greater challenges to computational methods (Wen et al. 2018); for example, almost opposite conclusions were drawn from two methodologically different studies of the same assemblage of *Xiphophorus* swordtail fish (Cui et al. 2013; Kang et al. 2013). The challenge of characterising introgression in adaptive radiations remains open and requires both taxonomic completeness and sophisticated methodological approaches.

The charismatic Neotropical *Heliconius* butterflies have provided some of the most compelling examples of interspecific gene flow occurring pervasively across the genome and at specific adaptive genes (Table 1). The natural propensity of *Heliconius* and the sister genus *Eueides* to produce hybrids in the wild (Mallet et al. 2007; Dasmahapatra et al. 2007) has generated an early interest in the genetic porosity of the species barriers (Beltrán et al. 2002; Bull et al. 2006). Further in-depth studies revealed that ecologically divergent species in the *melpomene-cydno*-silvaniform clade (MCS) can share variation at up to 40% of the genome in sympatry (Kronforst et al. 2006; Nadeau et al. 2012; Nadeau et al. 2013; Mérot et al. 2013; Kronforst et al. 2013; Martin et al. 2013; Martin et al. 2018). Loci responsible for aposematic and mimetic wing patterns are especially likely to be shared between species (Heliconius Genome Consortium 2012; Pardo-Diaz et al. 2012; Enciso et al. 2017; Jay et al. 2018), providing a source of genetic variation in a strongly selected trait *Heliconius heurippa*, also belonging to the MCS clade, remains the best documented case of homoploid hybrid speciation (Salazar et al. 2010; Schumer et al. 2014), in which the adaptive red wing pattern introgressed from *H. melpomene* into the *H. cydno* background contributes to reproductive isolation between species (Mavarez et al. 2006). Introgression at the *cortex* locus between *H. cydno* and *H. melpomene* facilitated the occupation of atypical mimetic niches in both species (Enciso et al. 2017). More surprisingly, the complex *Dennis/Ray* pattern found in *H. melpomene* and relatives, as well as in *H. elevatus*, is a product of bidirectional adaptive introgression between lineages up to 5 million years apart (Heliconius Genome Consortium 2012; Wallbank et al. 2016). Similarly, introgression of an inversion fixed in *H. pardalinus* triggered the evolution of polymorphic wing patterns in *H. numata* (Jay et al. 2018).

**Table 1.** Loci and direction of introgression varies between species. An overview of incongruences at the colour pattern loci detected by inspecting ML gene trees, and genome-wide admixture found with TreeMix and MPL networks. Frequency of the clusters counted among autosomal gene trees. Novel findings highlighted in yellow.

| Recipient | B/D Donor | Yb/Cr | Ac/Sd | Ro | K | frequency | Autosomes-wide |
| --- | --- | --- | --- | --- | --- | --- | --- |
| *H. hecalesia* | *H. clysonymus*/<br>*H. hortense* | *H. clysonymus*/<br>*H. hortense* | *H. clysonymus*/<br>*H. hortense* | *H. clysonymus*/<br>*H. hortense* |  | 0.140 | Variation shared with *H. clysonymus*,<br>*H. sapho* and *H. erato*<br>clades. |
| *H. numata* |  |  |  |  |  | 0.003 | *H. melpomene* E |
| *H. hecale clearei* | *H. pardalinus*; *H. numata*; *H. ethilla* |  |  |  |  | 0.046; 0.005;<br>0.019 |  |
| *H. elevatus* | *H. melpomene* E |  | *H. melpomene* E,<br>*H. cydno* |  |  | 0.003; 0.001 |  |
| *H. melpomene* E | *H. elevatus* |  |  |  |  | 0.003 |  |
| *H. pardalinus/elevatus* |  | *H. melpomene*/<br>*H. cydno* |  |  |  | 0.000; 0.004 | *H. melpomene* E |
| *H. timareta/heurippa* | *H. melpomene* E | *H. melpomene* E |  | *H. melpomene* E | *H. melpomene* E | 0.043 | *H. melpomene* E |
| *H. timareta* |  |  | *H. melpomene* E |  |  | 0.090 |  |
| *H. heurippa* | *H. melpomene* W |  | *H. cydno*/*H. pachinus* |  |  | 0.008; 0.043 |  |
| *H. cydno/timareta* |  | *H. melpomene*<br>W/FG |  |  |  | 0.000 |  |
| *H. cydno/pachinus* |  | *H. melpomene* W |  |  |  | 0.043 | *H. melpomene* W |
| *H. m. malleti* Colombia |  |  | *H. cydno*/*H. timareta* |  |  | 0.000 |  |
| *H. pardalinus/elevatus*/<br>*hecale/atthis/ethilla* |  |  |  |  |  | 0.008 | *H. melpomene* E |

It is virtually unknown whether the phenomena documented in the relatively recently emerged (4.5-3.5MYA) (Kozak et al. 2015) MCS clade are typical of the genus as a whole. Museum collection data suggest that hybridisation happens most frequently among the MCS species (Mallet et al. 2007), which can also be hybridised in captivity (Gilbert 2003). However, many hybrids are also known between *H. erato* and *H. himera*, as well as within the genus *Eueides* (Mallet et al. 2007). Here we generate alignments for 7324 orthologous, protein-coding genes obtained from a comprehensive whole-genome resequencing data set of 40 of the 45 recognized *Heliconius* species, six of the 12 *Euides* species (the most closely related genus) and two outgroup species to investigate the prevalence of hybridisation in *Heliconius*, quantify its extent, and compare the processes producing discordance at several loci. We demonstrate varied amounts of phylogenetic incongruence across the genus, related to heterogenous levels of interspecific gene flow. Instances of hybridisation at adaptive loci and more broadly across the genome, are found across the radiation, although they are far more frequent in the relatives of *H. melpomene*.

Our study dissects the evolution of two genera across a continent to demonstrate that gene flow can be a ubiquitous and significant force in shaping an adaptive radiation, and to provide first estimates of the probability of such gene flow and adaptive introgression. We show that a misleadingly well-supported and resolved tree can be recovered despite incongruence, and that previously unknown, complex hybridisation event can thus be missed. We propose that phylogenetic networks should become *de rigour* tools in the study of adaptive radiations.

## RESULTS AND DISCUSSION

### A genomic tree for the heliconiines

We first constructed a bifurcating phylogeny for our genome-wide data. Mapping genome-wide short read data from 144 individuals in 45 species (S1 Table) to the *H. melpomene* reference did not result in full coverage of the distantly related species (S2 Table), but we were able to recover 7264 high quality, orthologous CDS alignments with less than 4% missing data (S3 Table). Filtering for autosomal exome sites with biallelic, non-singleton SNPs without missing data produced a 122,913 bp supermatrix. This matrix generated a Maximum Likelihood tree that is resolved with full bootstrap support (S1 Figure), except for uncertain placement of the *H. telesiphe/hortense/clysonymus* clade (bootstrap support 62/100). This concatenation tree is not substantially different from the results obtained by either ASTRAL or MP-EST multispecies coalescent methods (Fig. 1), which model the species tree from the distribution of gene trees. Both MSC methods also yield well resolved phylogenies and, in the case of ASTRAL, completely supported topologies (Fig. 1). The MSC trees differ from each other at only two relatively recent splits, while the concatenation phylogeny differs from ASTRAL at three out of 56 (Fig. 1).

**Figure 1.**
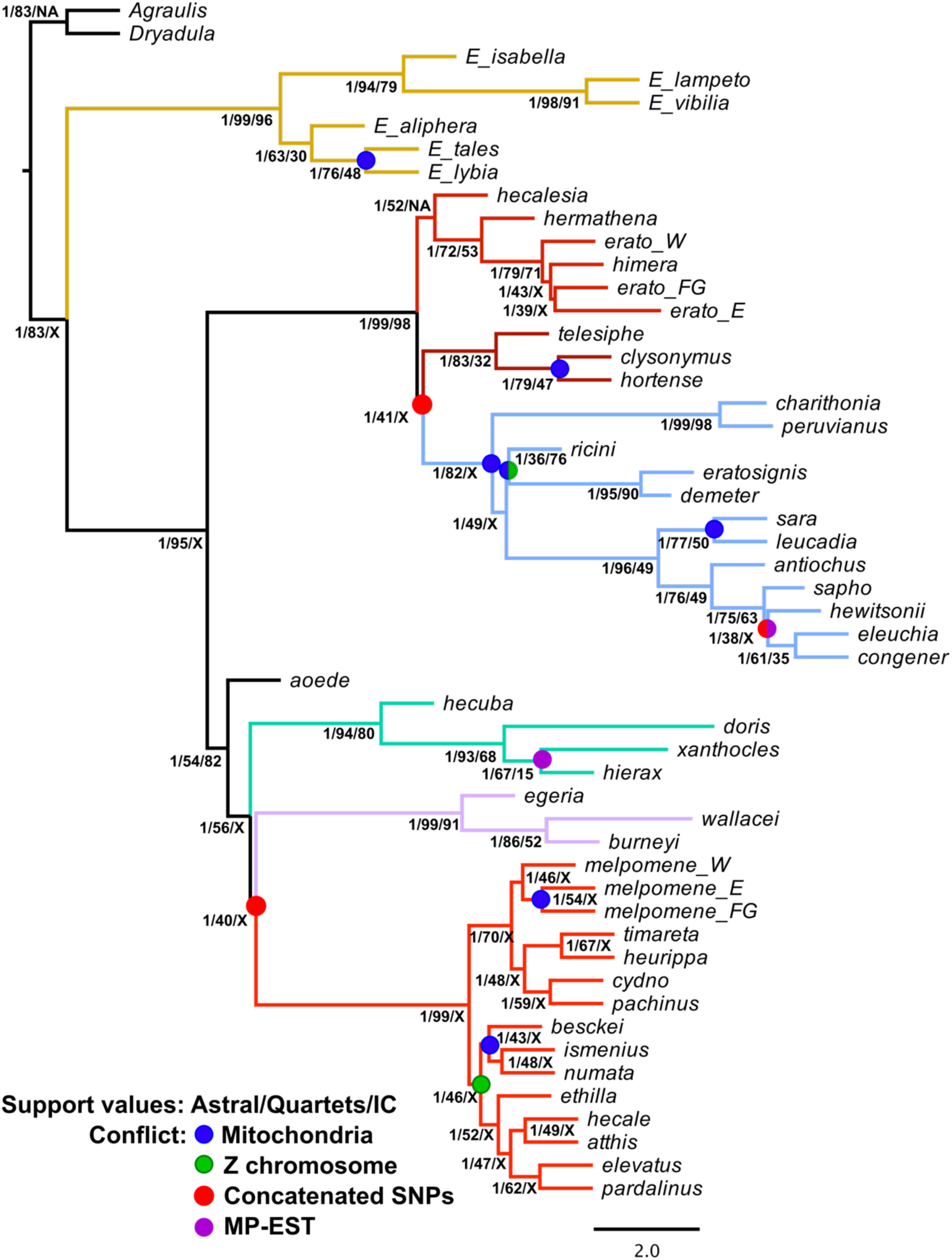
Well-resolved phylogeny of Heliconiini obscures the underlying incongruence. The topology identified by the multispecies coalescent method ASTRAL is well supported, although many of the nodes are not represented in several of the gene trees (numerical values). Topologies are consistent between coalescent and concatenation methods based on whole genome data, but conflicted mainly with the mitochondrial phylogenies (blue dots). Branch colours correspond to previously defined clades (Brown 1981; Kozak et al. 2015): red – *melpomene*/Silvaniforms; violet – *egeria*; green – *hecuba*; blue – *sara*; crimson – *clysonymus*; scarlet – *erato*; brown – *Eueides*.

At one level, these well resolved trees clarified some uncertainties from previous work (Beltran et al. 2007; Kozak et al. 2015), including placement of the *H. hecuba* and *H. egeria* groups, relations in the *H. sapho* clade, and the position of *H. besckei*. Nonetheless, similar to other large phylogenetic studies (Brawand et al. 2014; Jarvis et al. 2014; Fontaine et al. 2014), none of the individual gene trees showed exactly the same topology as the species tree. making us realise that the well-supported bifurcating trees do not fully represent the underlying signal in the genomes within this clade.

### Genome-wide phylogenetic incongruence

Rapid radiations of closely related species are likely to show considerable phylogenetic incongruence due to incomplete lineage sorting and introgressive hybridization. This is especially likely to be the case in *Heliconius*, where population sizes are large (Flanagan et al. 2010), speciation events are clustered in time (Kozak et al. 2015), inter-specific hybrids are found in the wild (Mallet et al. 2007), and there is strong evidence for extensive introgression of closely related sympatric species (Martin et al. 2013; Martin et al. 2018). Indeed, despite the apparently well-supported phylogeny, we found considerable phylogenetic incongruence across the genome in our data. For example, Robinson-Foulds pairwise distance was 0.745 out of 1.0 for the 6848 autosomal phylogenies and 0.699 for the 416 loci on the sex-linked Z chromosome, indicating that any two gene trees are likely to contain multiple differing partitions. Among the 56 nodes between species and major subspecies, less than a half (26) are resolved in an autosomal Majority Rule Consensus tree (S2-S4 Figures). The relative tree certainty (TCA) of the consensus is 0.322 on a 0-1 scale, whereas many supported branches score low on the Information Criteria (IC/ICA), indicating that there is one (IC) or multiple (ICA) alternative groupings found at high frequency among the gene trees (Salichos & Rokas 2013) (S2 Figure). Brower and Garzon-Orduña (2017) suggested that the phylogenetic incongruence among Heliconiini is an artifact of missing data. This is unlikely, as the 7324 nucleotide matrices recovered here are nearly complete (>96%) and only 0.87% of the sites were removed as incomplete or ambiguous (S3 Table). Phylogenetic studies using modern statistical methods are typically robust to far higher levels of missing data (Wiens & Morrill 2011; Roure et al. 2013).

It has been suggested that sex-linked markers might be more reliable than autosomes for phylogenetic reconstruction, due to faster coalescence at lower effective population size (3/4 of the autosomal *N*_*e*_ at the Z), and reduced likelihood of hybridisation as a result of Haldane’s Rule (Zhang et al. 2013; Fontaine et al. 2014). We found that there is indeed significantly greater concordance between species tree and the Z chromosome trees, as compared to autosomal trees (gene tree-species tree triplet distance: 10.3*10^6^ *vs* 12.5*10^6^, Wilcoxon’s test *p*=3*10^−11^) (S3 Figure). Notably, many nodes within the MCS clade are resolved among Z chromosome trees, and the *melpomene/cydno* clade is monophyletic. This agrees with genomic studies of house mouse subspecies (White et al. 2009), fruit flies (Garrigan et al. 2012; Pease & Hahn 2013) and *Anopheles* mosquitoes (Lee et al. 2013; Fontaine et al. 2014) showing deeper coalescence and lower levels of phylogenetic discordance at the X chromosomes. A comparison of gene tree discordance levels, and coalescent trees between autosomes and the Z demonstrates that the sex chromosome in *Heliconius* is likely subject to slightly lower levels of gene flow (S4 Figure). Nonetheless, the Z-linked consensus did not resolve the relative positions of the major clades unequivocally (S2-S3 Figures) and there were roughly similar levels of conflict with the species tree between sex-linked and autosomal loci (S5 Figure). Thus, even the Z chromosome yields inconsistent the phylogenic estimates (S4 Figure). This may be a result of an ancient introgression of the entire chromsome, since with our larger species sampling we confirm previous demonstration of Z chromosome incongruity between the MCS clade and the Silvaniform group containing *H. ethilla, H. hecale* and *H. pardalinus* (S6 Figure) (Zhang et al. 2016).

Similarly, the whole mitochondrial phylogeny was strongly conflicted with the multispecies coalescent trees (Fig. 1), demonstrating that organellar history is unlikely to be a good proxy for phylogeny at the species level (Hurst & Jiggins 2005). Generally, the same branches tended to be short and unsupported in concatenation and coalescent analyses of mitochondrion-, Z-and autosome-linked markers (Fig. 1, S1-S3 Figure, S6 Figure). This corroborates the idea that some of the diversification events among Heliconiini occurred nearly simultaneously and are unlikely to be resolved in a bifurcating tree.

### Hybridisation between species has been common throughout the radiation of *Heliconius*

We next sought to document the extent and degree of discordance across the phylogeny, and distinguish between incomplete lineage sorting and gene flow. We did this in two ways, first modeling the extent of hybridization in the history of the radiation in the software PhyloNet, using the Maximum Pseudo-Likelihood network algorithm for distinguishing admixture from Incomplete Lineage Sorting based on gene tree topologies and branch lengths (Yu & Nakhleh 2015). Here, an inference of speciation history networks from 6848 gene trees revealed a pattern of gene sharing within all major clades of the radiation (Fig. 2). This was particularly true within the *H. melpomene, H. cydno* and Silvaniform clade (MCS), where our approach supports previous work documenting extensive admixture, including the hybrid speciation of *Heliconius heurippa* (Mavarez et al. 2006; Salazar et al. 2008; Salazar et a. 2010), admixture between *H. cydno/timareta* and subspecies of *H. melpomene* (Nadeau et al. 2013; Martin et al. 2013; Enciso et al. 2017), and the exchange between *H. melpomene* and *H. elevatus* (Heliconius Genome Consortium 2012; Wallbank et al. 2016). This result made us confident that outcomes of this analysis can be trusted.

**Figure 2.**
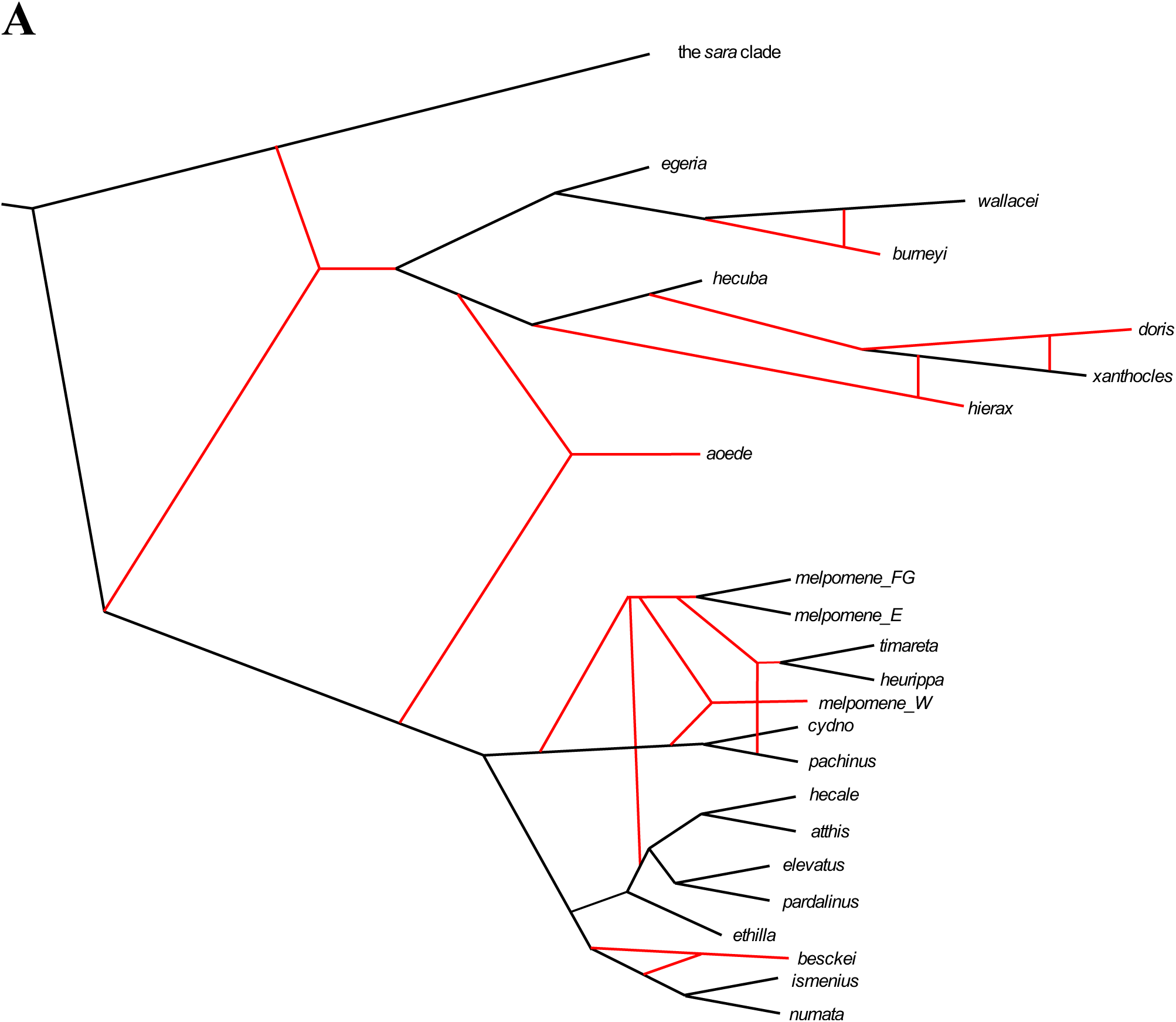
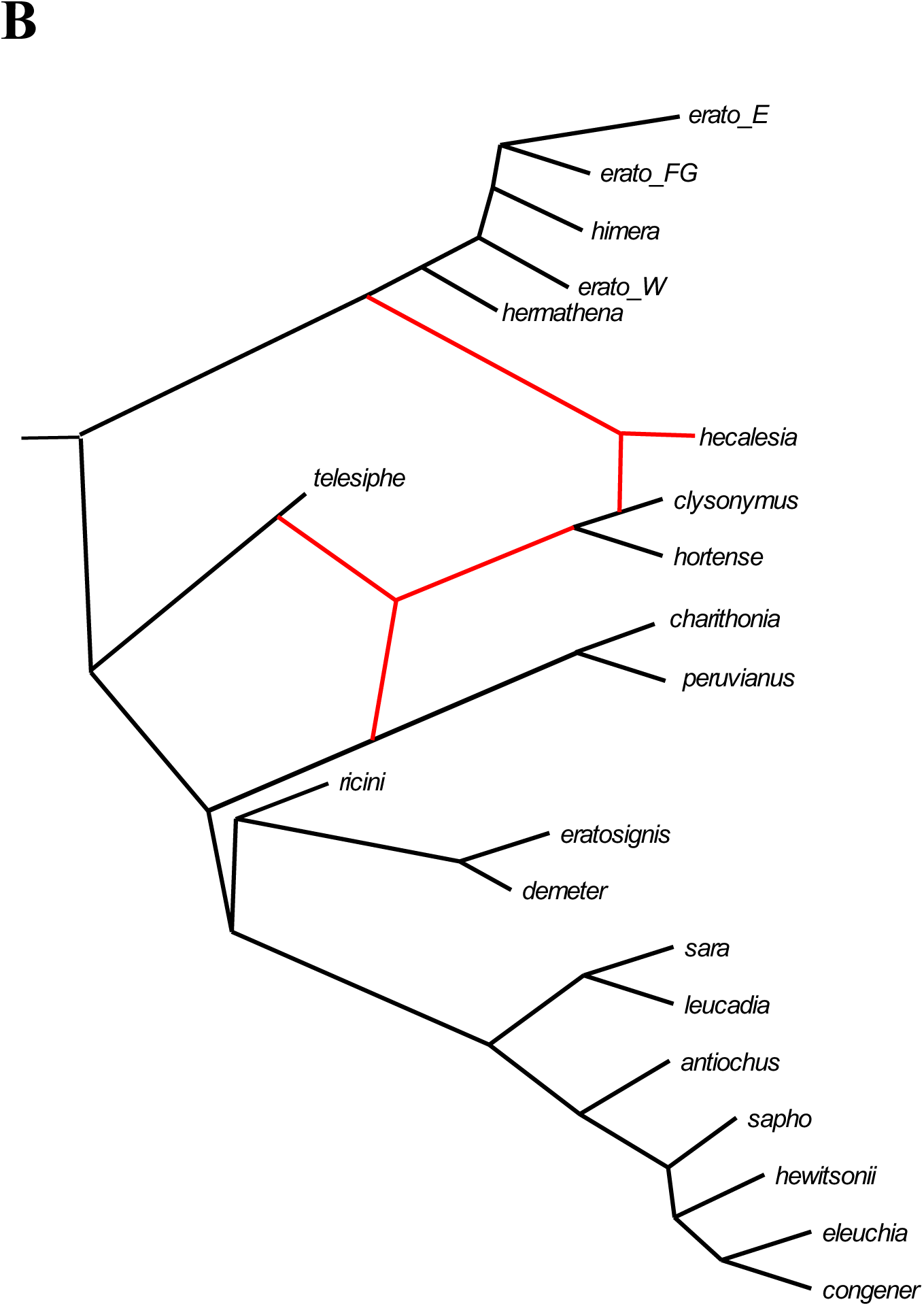
Evidence of introgression is found across the entire *Heliconius* radiation. Networks inferred under Maximum-Pseudolikelihood (MPL) based on 6724 autosomal ML gene trees distinguish between introgression and incomplete lineage sorting, revealing previously unknown admixture events (red edges). (A) The analysis of *H. melpomene* and cognates reveals several unknown introgressions closer to the root of the tree. Known events are recapitulated, demonstrating robustness of the approach. (B) Relatively few admixtures occurred in the evolution of the *Heliconius erato/sara* clade, but *H. hecalesia* may be a previously unknown recent hybrid.

In the small clades containing *H. egeria, H. hecuba* and *H. aoede*, we found similar evidence for multiple instances of admixture, particularly between all species in the *H. hecuba* quartet (Fig. 2B). Gene flow is also apparent between the ancestors of modern clades, including a potential hybrid origin of *H. aoede*. This finding explain the unusual combinations of morphological and ecological traits in these species, which used to be classified in separate genera (Brown et al. 1981; Brower 1994) prior to molecular evidence (Beltran et al. 2007; Kozak et al. 2015). Finally, in the large group of *H. erato* (*sensu* Brown 1981)we detected admixtures between the triplet of (*H. telesiphe*,(*clysonymus, hortense*)) and both *H. erato* and *H. sara* clades (Fig. 2A). Notably, there are fewer instances of reticulation than in all the other groups consistent with other evidence (Fig. 4, S9 Figure). The most likely network estimated for the genus *Eueides* contains an unusual reticulation pattern leading to none of the included taxa (S1 Figure). We suspected that this pattern may be a sampling artefact, as only half of the species are included with one individual each, and divergence from the reference genome makes the *Eueides* alignments less complete and reliable.

Although MPL networks are currently the most robust and computationally efficient tools to model gene flow in face of ILS (Wen et al. 2018), it is unknown if the estimates are sensitive to errors in gene tree estimation and missing data as in case of *Eueides*. To account for this possibility we also used the Ancestral Recombination Graph algorithm implemented in TreeMix (Pickrell & Pritchard 2012), which analyses genome-wide variation under a Gaussian drift model, interpreting deviations from expected differentiation as evidence for gene flow. When applied to the SNP supermatrix, TreeMix again detects known events in the MCS clade, confirming the robustness of the technique, even when applied at the relatively deep level of a genus (Fig 3).

**Figure 3.**
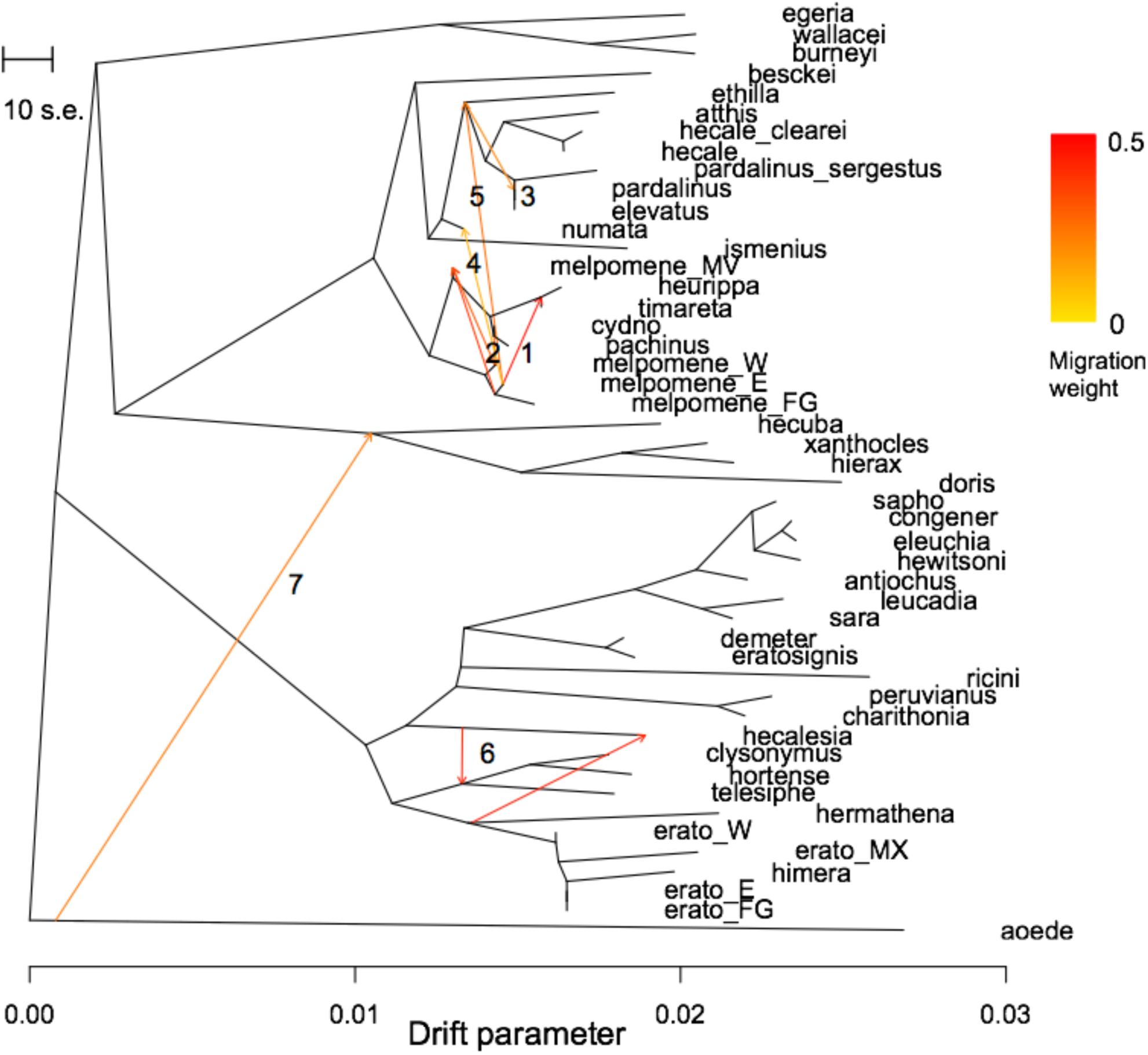
The extent of interspecific gene flow varies across the tree. TreeMix inference of splits and mixture from autosomal SNPs. Migration edges (1-7) are inferred on a phylogenetic tree built from allele frequencies under a Gaussian genetic drift approximation. Colours of the edges correspond to the proportion of the genome exchanged.

The ARG shows gene flow from *H. hecalesia* into the *telesiphe* clade, and from the *erato* lineage into *H. hecalesia* (Fig. 3). This suggests a hybrid origin for *Heliconius hecalesia* (Fig. 2A). In contrast, and somewhat surprisingly, we find no evidence for a hybrid origin of *H. hermathena*. This species, found in the white sand forests of Brazil, has a wing pattern resembling a combination of the zebra-patterned of *H. charithonia* and the red-banded *H. erato hydara* (Brown & Benson 1977; Mavárez & Linares 2008; Mavárez et al. 2006).

In the small clades including *H. egeria, H. hecuba* and *H. aoede*, we find evidence for pervasive admixture, particularly between all species in the *hecuba* quartet (Fig. 2B). In contrast, TreeMix detects only a single gene flow event from *H. aoede* into the *egeria/hecuba* clade (Fig. 3). Gene flow is also apparent between the ancestors of modern subgenera, including a potential hybrid origin of *H. aoede*, which may explain the unusual trait combinations in these species that were sometimes classified in separate genera (Brown et al. 1981; Beltran et al. 2007; Kozak et al. 2015) (Fig. 2, 3). The Treemix ARG and PhyloNet network approaches do not uncover an exactly identical set of introgression results, as the ARG does not detect the exchanges in and between the small *H. egeria* and *H. hecuba* clades, or some of the events previously documented between species of the Silvaniform group (Jay et al. 2018) and *H. melpomene* (Zhang et al. 2016). The discrepancies are expected between two widely different techniques, and the Treemix algorithm assumes that the underlying sequence of events was largely tree-like (Pickrell and Pritchard 2012). Although Treemix is designed for populations and less commonly used to examine divergent species (but see Zhan et al. 2014), it infers a basic tree similar to that found by the phylogenetic approaches that do not model introgression (Fig. 1).

Distributions of allele frequencies show that the extent of introgression between species varies between clades within *Heliconius*. When all possible combinations of taxa are examined, most of the quartets in the MCS clade (22 species, 3309790 SNPs) show some admixture (*f4*<0; *Z*-score<-4.47 after Bonferroni correction for the number of lineages) (Fig 4). In contrast, most *f4* values computed from the *H. erato* group (20 species, 3152880 SNPs) are not significant and thus indicate a relatively better fit to a simple bifurcating tree (Peter 2016). This summary statistic allows a crude comparison of levels of admixture across the genus, but it remains computationally difficult to examine gene flow between more than a handful of taxa using these approaches (Martin et al. 2015; Pease & Hahn 2015).

**Figure 4.**
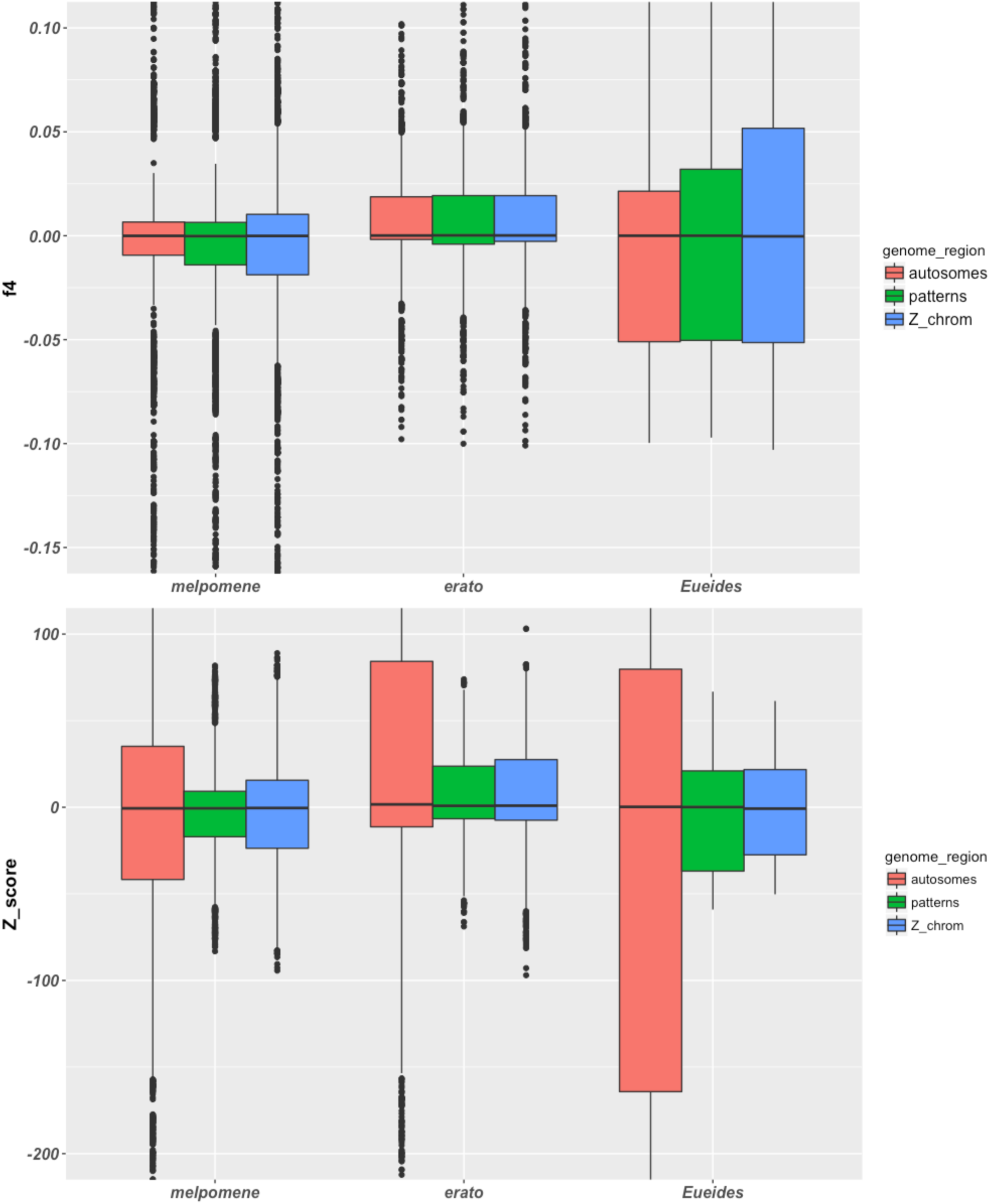
Structure of the *Heliconius erato* clade is the closest to the bifurcating tree model. (A) When introgression among all possible quartets is tested in the major clades, more admixture events (negative *f4* statistics) are found in the *H. melpomene* than *H. erato* group. *Eueides* shows high variation possibly due to incomplete sampling. Gene flow is more common at putatively neutral autosomal sites than adaptive patterning or sex-linked loci. (B) *Z*-scores for all possible *f4* estimates in each clade calculated by jackknifing. Values below −4.47 indicate a significant test at the 0.01 level after correction for multiple testing.

### Adaptive introgression at the wing pattern loci

A characteristic aspect of *Heliconius* evolution is the adaptive introgression of five colour pattern loci, corresponding to a small number of protein-coding genes (Table 2) (reviewed in Kronforst and Papa 2015). For many of these genes more recent work has identified intervals that are associated with specific aspects of color and pattern that are thought to contain regulatory regions responsible for differential expression (Table 2). Introgression driven by selection for Mullerian mimicry has been documented extensively (Salazar et al. 2010; Pardo-Diaz et al. 2012; Heliconius Genome Consortium 2012; Wallbank et al. 2016; Zhang et al. 2016; Jay et al. 2017), but only in the *H. melpomene* and Silvaniform (MCS) clade of *Heliconius*, which includes less than a third of all species. By comparing the topologies at the colour pattern loci for nearly all species in the genus against the overall species tree, we provide a uniform framework in which to gauge the amount of introgression across around these functionally important regions. Topologies around color pattern loci differed from the species tree (*p*<0.001, SH test) and in many cases, primarily at the *optix* and *cortex* loci, showed multiple departures from the expected topology (Table 1). The majority of the differences are found in the MCS clade, where introgression reaches considerable complexity across genomic and geographic regions (Wallbank et al. 2016; Enciso et al. 2017). Again, we verify our approach by recovering the known signals the *optix* and *cortex* loci (Table 1), but it is the first time that introgression is demonstrated in other genomic intervals.

**Table 2.**
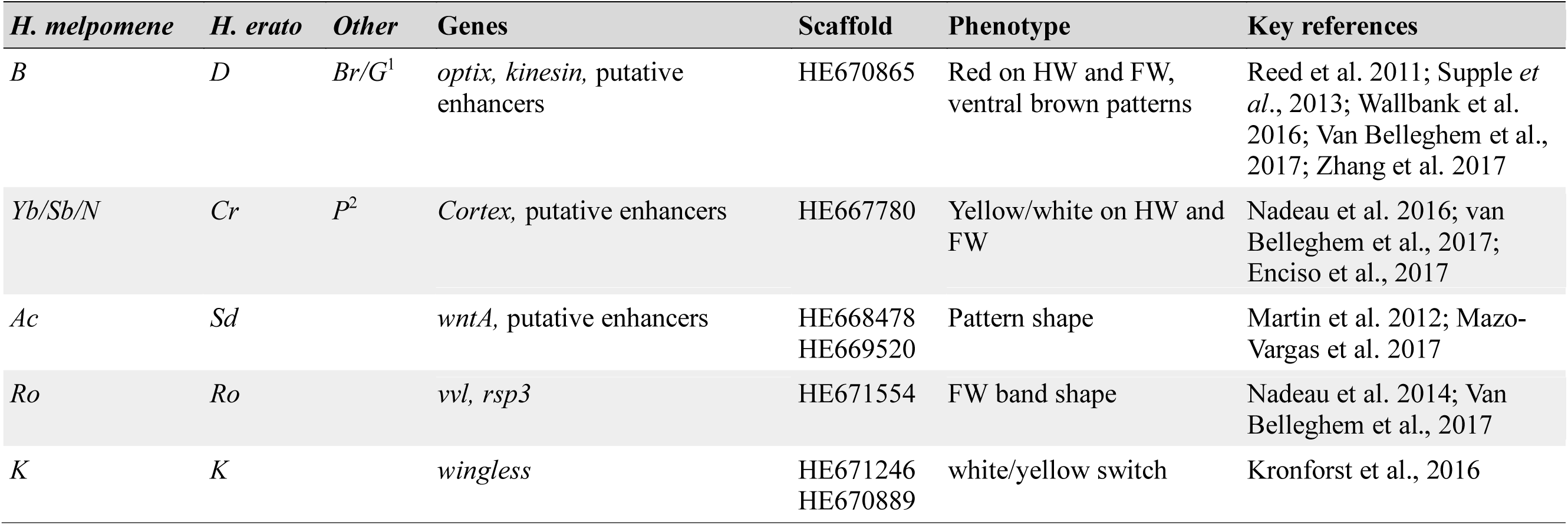
Major wing pattern and colour loci in *Heliconius*. The key loci are named differently in divergent species (Sheppard et al. 1985), but in all cases the approximate genomic location has been identified, good candidate genes have been identified and intervals containing functional variation have been localized (Reed et al., 2011; Martin et al., 2012; Nadeau et al. 2016; Wallbank et al., 2016; Van Belleghem et al. 2017). Scaffold numbers refer to the *Hmel v1* assembly. ^1^Brown patterns in *H. cydno* and *H. pachinus* (Chamberlain et al. 2011); ^2^The *Pushmipullyu* supergene controlling most of the wing patterning in *H. numata* (Joron et al. 2011). HW: hindwing; FW: forewing.

In contrast, the *H. hecalesia/clysonymus/hortense* is the only case of wing pattern introgression found among the 19 species of the *H. sara/erato/telesiphe* subgenus. No pattern introgression is found among the three smaller clades (for example Fig. 6), despite evidence for gene flow in other regions (Fig. 2, 3). Although we cannot rule out an influence of sampling bias, as substantially more individuals have been sequenced in the MCS group, our data nonetheless indicate that introgression around color pattern loci is more pervasive in this clade.

**Figure 6.**
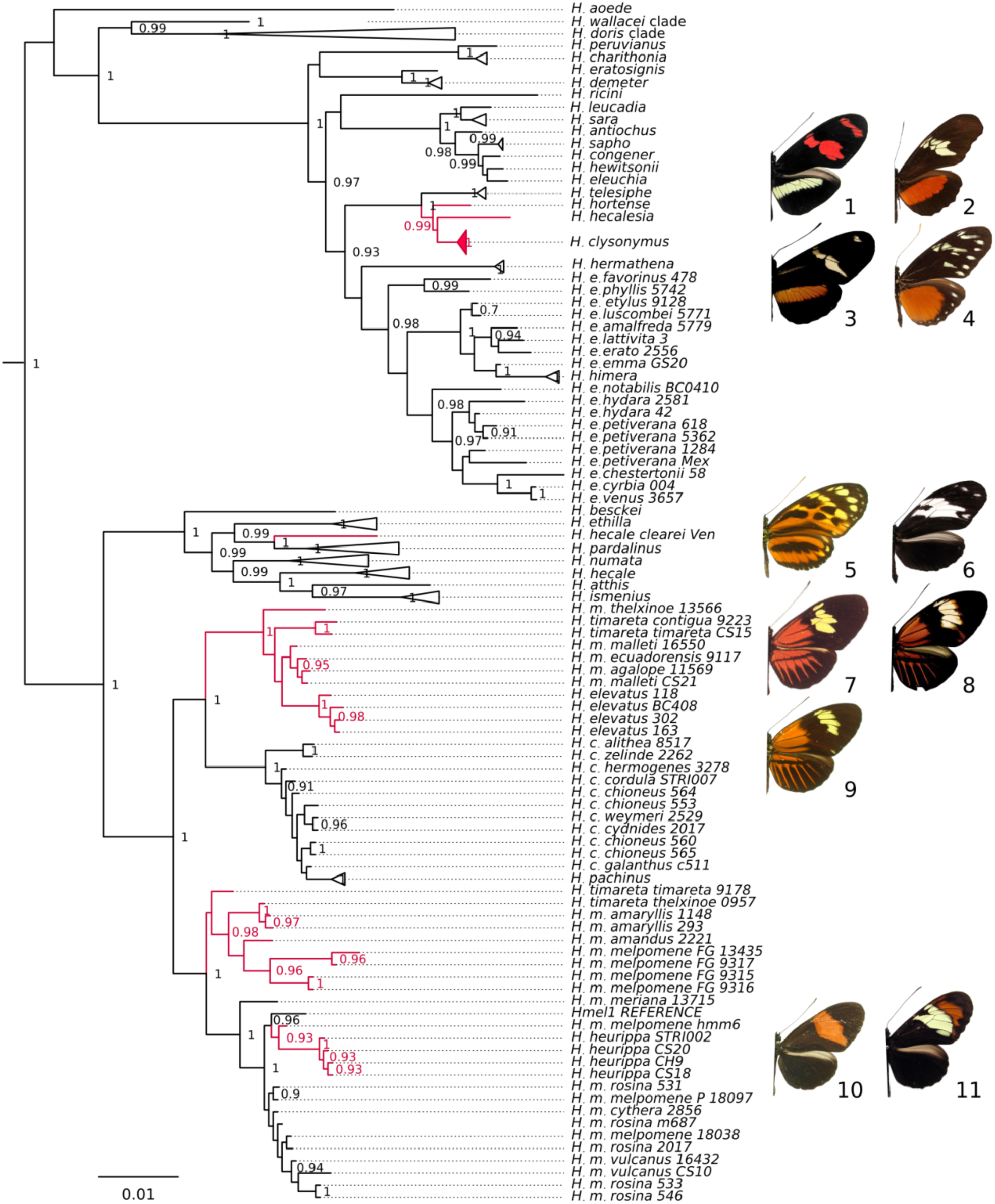
Pervasive introgression across the species boundary at the red patterning locus. Branches in unexpected positions labeled red. The ML tree was reconstructed for the 360,000-380,000 bp region on the B/D scaffold, including the main peaks of association with patterns in *H. erato* and *H. melpomene*. Intraspecific relations for species not discussed in text are collapsed. Outgroups and parametric support values <0.9 not shown. 1. *H. telesiphe sotericus,* 2. *H. hortense*, 3. *H. clysonymus hygiana*, 4. *H. hecalesia formosus*; 5. *H. hecale felix*, 6. *H. hecale clearei*; 7. *H. timareta timareta*, 8. *H. melpomene malleti*, 9. *H. elevatus*; 10. *H. melpomene melpomene* (Magdalena Valley), 11. *H. heurippa*.

We find new evidence for gene flow at the *cortex* locus (Nadeau et al. 2016) which is responsible for the diverse white and yellow patterns across the genus (Sheppard et al. 1985; Kronforst & Papa 2015; Nadeau et al. 2014). At a region previously highlighted (Nadeau et al. 2012) (310-330kbp in scaffold HE667780) *H. melpomene* and *H. timareta* alleles cluster with Silvaniforms (aLRT > 0.95); further downstream *H. melpomene, H. timareta* (570-620 kbp) and *H. cydno* (590-620 kbp) share alleles, as well as a cluster of sequences similar between *H. elevatus/pardalinus* and *H. melpomene/cydno* (570-600 kbp). This last association is also observed in the 670-690 kbp window containing *cortex* (Nadeau et al. 2014).

At the *WntA* locus, we again find topologies indicating Amazonian *H. melpomene* introgression into *H. timareta* and *H. elevatus*. In the *WntA* (Martin et al. 2012) windows (450-490 kbp), sequences of *H. heurippa* cluster with *H. cydno*, upholding the view that the former arose by hybrid speciation from a yellow-patterned race of *H. cydno/timareta*.

Most of the variation in the *optix* region is consistent with the genome-wide lack of resolution in the *H. melpomene/cydno*/Silvaniform clade and confirms known events. The greatest number of discordant branches are among the *H. melpomene/cydno* clade at 360 to 380 kbp (Fig. 6), the section controlling both *H. melpomene* (Wallbank et al. 2016) and *H. erato* red ray patterns (Supple et al. 2013; van Belleghem et al. 2017). Intriguingly, alleles from *H. hecale clearei* cluster with the *H. pardalinus/H. elevatus* sequences in eight windows (Fig. 6), perhaps related to the complete loss of orange patterning in this uniquely black and white Silvaniform.

*Heliconius* stand out as having multiple examples of introgression of unlinked loci enabling the rapid evolutionary development of complex patterns, which comprise a patchwork of elements sometimes derived from different sources (Table 1)(Wallbank et al. 2016). A large number of genomic studies of interspecific gene flow have found introgressions of small genome regions driven by natural selection for critical adaptations, such as the hypoxia resistance *EPAS1* haplotype (Denisovans → anatomically modern Tibetans) (Huerta-Sánchez et al. 2014), the *Vgsc-1014F* insecticide-resistance mutation (*Anopheles gambiae* → *A. colluzzi*) (Clarkson et al. 2014; Norris et al. 2015), or the *ALX1* alleles determining diverse beak shapes among Darwin’s finches (*Geospiza*) (Lamichhaney et al. 2015). Whereas adaptive introgression has been localised to a single part of the genome in all but one other animals studied to date (but see Stryjewski & Sorenson 2017), it is clear that *Heliconius* wing patterns involve introgressions at multiple loci and between different combinations of species. For instance, *H. elevatus* derives *WntA* and *optix* alleles (Table 1) (Wallbank et al. 2016) from Amazonian *H. melpomene*, but some of the *WntA* haplotype appears to be transmitted from *H. cydno*, possibly *via H. melpomene*. Similarly, fragments of the *optix, WntA* and *cortex* sequences in *H. elevatus* are derived from either *H. melpomene* or *H. cydno*.

### Mosaic genome in *Heliconius hecalesia*

We identify the first cases of gene flow among distantly related species in the *H. erato* clade, demonstrating that admixture has not been limited to the well-studied MCS group. We infer two instances of gene flow. into both the ancestor of *H. hortense* and *H. clysonymus,* and into *H. hecalesia*. The position of the (*H. telesiphe*,(*hortense,clysonymus*)) triplet (THC) is the only unsupported branch in the genome-wide phylogeny (Fig. 1; S2 Figure), and no specific placement is found in a majority of gene trees (IC=0; S3 Figure). This uncertainty is not a result of coalescent stochasticity, as both MPL networks and TreeMix account for ILS and still recover signals of admixture from both the clades of *H. erato* and *H. sara* into THC (Fig. 2A, 3). *H. hecalesia* latter has especially complex ancestry, and although usually grouped with *H. erato* in simple trees (Fig. 1) (Kozak et al. 2015), appears to be nearly equally diverged from *H. erato, H. telesiphe* and *H. sara* (Fig. 5). Results of network modelling are corroborated by the *D* statistic of admixture (Durand et al. 2011), which is highly positive and statistically significant for all tests where *H. hecalesia* is the recipient of admixture from the *H. clysonymus* and *H. sara* clades (Table 3). However, consistent with the pattern of genome-wide variation, there is evidence for stronger gene flow between similarly patterned *H. hecalesia* and *H. clysonymus* (*D*=0.35; *p*<0.0001) or *H. hortense* (*D*=0.38; *p*<0.0001) than the dissimilar *H. sara* (*D*=0.17). At the higher phylogenetic level there is also evidence for gene flow between the THC clade and *H. hecalesia*, but not *H. erato* (Table 3), raising the possibility that admixture took place between the ancestors of modern clades, leading to incongruence deep in the tree (S1 Figure).

**Table 3.** D-statistic values (ABBA/BABA tests) of admixture between *H. hecalesia* and relatives. Lower section of the table describes tests treating entire clades as taxa, to detect ancient admixture. P2 is the hypothetical recipient and P3 the donor of variants. Positive *D* values are evidence for admixture after accounting for ILS. *p*-values calculated by block jacknifing.

| *P1* | *P2* | *P3* | *D* | error(*D*) | *p*-value |
| --- | --- | --- | --- | --- | --- |
| *erato* FG | *hecalesia* | *clysonymus* | 0.349 | 0.009 | <0.0001 |
| *erato* FG | *hecalesia* | *hortense* | 0.383 | 0.009 | <0.0001 |
| *erato* FG | *hecalesia* | *telesiphe* | 0.269 | 0.008 | <0.0001 |
| *erato* FG | *hecalesia* | *charithonia*+*peruvianus* | 0.208 | 0.005 | <0.0001 |
| *erato* FG | *hecalesia* | *sara*+*leucadia* | 0.170 | 0.005 | <0.0001 |
| *erato* East | *hecalesia* | *clysonymus* | 0.354 | 0.009 | <0.0001 |
| *erato* East | *hecalesia* | *sara*+*leucadia* | 0.178 | 0.004 | <0.0001 |
| *erato* clade | *hecalesia* | *clysonymus* clade | 0.305 | 0.008 | <0.0001 |
| *erato* clade | *hecalesia* | *sara* clade | 0.171 | 0.004 | <0.0001 |
| *hecalesia* | *erato* clade | *clysonymus* clade | -0.305 | 0.008 | *NA* |
| *hecalesia* | *erato* clade | *sara* clade | -0.171 | 0.004 | *NA* |

**Figure 5.**
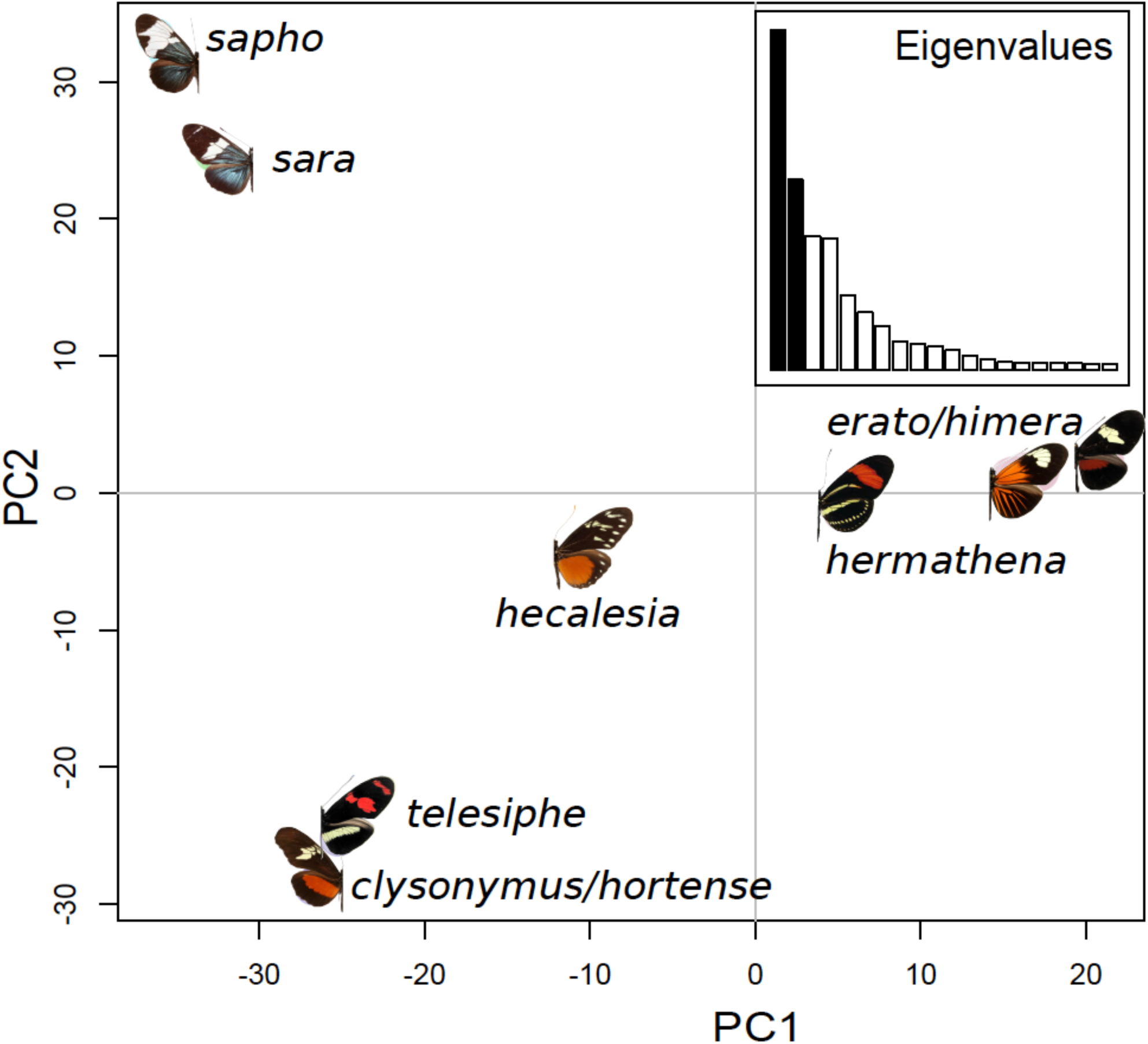
Ambiguous genomic composition of *H. hecalesia*. Although usually recovered as the sister species of *H. hermathena* and *H. erato* in bifurcating phylogenies, H. hecalesia shares substantial amount of variation with two other clades. Principal Component Analysis of variation in the autosomal exonic SNPs within the *H. erato/sapho* clade. First two PCs plotted, accounting for over half of the variation.

For the first time we find evidence outside of the MSC clade for introgression of multiple colour pattern alleles, from *H. hecalesia* into the sympatric THC lineage, possibly due to selective pressure on mimetic appearance. *H. clysonymus* and *H. hortense* display the unusually wide red/orange bar on the hindwing, absent from their relative *H. telesiphe*, but shared with the distantly related *H. hecalesia* (Fig. 5). Based on the phenotype we predict that, at the key wing patterning loci (Table 2), sequences from these three species should cluster together to the exclusion of *H. telesiphe*. Sequences from the three species indeed cluster exactly at the 360-380kbp interval of scaffold HE670865 (aLRT>0.95; *p*<0.001, SH test), which aligns to the specific region of the *D* locus controlling the red pattern in *H. erato* (Supple et al. 2013) (Fig. 6). This indicates that the alleles governing the red pattern in the three species are more similar than expected from the genome-wide phylogenies. As predicted if the sequences were introgressed after species divergence, the alleles from the three phenotypically similar butterflies appear younger (2.3-2.1 MA) in RelTime estimates than those of *H. telesiphe* (2.9MA), and much more recent than the original split between *H. hecalesia* and the THC clade (6.28-4.18 MA) (Kozak et al. 2015).

Whereas *optix* colours the band red (Zhang et al. 2017), the shape of bands is typically regulated by *WntA* (Mazo-Vargas et al. 2017), as well as by the *Ro* locus (Nadeau et al. 2014). As predicted from the shared phenotype of a wide hindwing band, we find that *H. hecalesia/clysonymus/hortense* cluster both at the *WntA* interval (HE667780: 450-490 kbp) (Martin et al. 2012), and in the section of *Ro* containing eight SNPs associated with pattern shape in *H. erato* (HE671554: 10-30kbp). Consistently, sequences of the phenotypically different *H. telesiphe* are divergent.

Adaptive gene flow between the three species is plausible, as *H. clysonymus* x *H. hecalesia* and *H. hortense* x *H. hecalesia* hybrids are known from the wild (Mallet et al. 2007), and *H. hecalesia* is sympatric with the other two species in parts of its range (Rosser et al. 2012). The subtle changes in the wing morphology of *H. hecalesia* across its range appear driven primarily by sexually-dimorphic mimicry of species in the Ithomiinae, with a wider hindwing band found further away from the Equator (Brown & Benson 1975). It seems likely that the precise mimicry between the distantly related *H. hecalesia* and *H. clysonymus/hortense* has resulted from localised introgression of regulatory loci in sympatry, similar to that seen between *H. melpomene/elevatus* (Wallbank et al. 2016).

### Neither concatenation nor coalescent trees adequately represent species history

There has been a marked shift over recent years away from phylogenetic methods that involve concatenation of data towards approaches that involve coalescent modelling. The proposed methods for inferring a species tree under the coalescent by modelling the incomplete sorting of loci represent a great improvement on the assumption that there is a common evolutionary history across all genome regions (Liu et al. 2010; Mirarab et al. 2014; Heled & Drummond 2010). Across the tree of life, from birds (Jarvis et al. 2014; Reddy et al. 2017) and mammals (Song et al. 2012) to fungi (Shen et al. 2016), treatment of individual gene trees under multispecies coalescent methods has yielded substantially different results to simple concatenation. However, both approaches still impose a bifurcating tree on the data and imply that speciation consists of a series of splits (Hahn & Nakhleh 2016). When applied to the adaptive radiation of *Heliconius*, both philosophies appear to produce results that fail to fully describe the evolutionary history of the species. Network modelling clearly demonstrates that introgression has happened throughout the evolution of the genus, and yet this process could easily be overlooked even with many state of the art phylogenetic methods. Appropriate reconstruction of speciation history needs to incorporate approaches that test for admixture more routinely (Long & Kubatko 2017; Pease & Hahn 2015). The main appeal of studying adaptive radiations is their power for analyzing trait evolution in a comparative framework, and a growing number of studies are looking at several *Heliconius* characters through this lens (Mallet et al. 2007; Briscoe et al. 2013; Rosser et al. 2015). Commonly, trait histories are best represented by exactly those regions that show high levels of introgression, and a comparative approach that used a single bifurcating species tree would give highly incomplete results. To accurately reflect the hidden uncertainties in the phylogeny of this exciting group, and to capture the potential of traits to be shared between species by genome-wide introgression, future work needs to utilize novel approaches that reflect the non-bifurcating reality of evolving genomes (Hahn & Nakhleh 2016; Bastide et al. 2017).

## METHODS

### Samples and DNA sequencing

Whole genomes of 11 species were re-sequenced for the first time: *Heliconius atthis, H. antiochus, H. egeria, H. leucadia, H. peruvianus, Eueides aliphera, E. lampeto, E. lineata, E. isabella, E. vibilia, Agraulis vanillae. H. antiochus* and *H. ricini* were collected in Bakhuis Mountains, Suriname. Other data were obtained from previous studies (Heliconius Genome Consortium 2012; Kronforst et al. 2013; Martin et al. 2013; Supple et al. 2013; Briscoe et al. 2013; Martin et al. 2016; Nadeau et al. 2016; Wallbank et al. 2016; Enciso et al. 2017; Jay et al. 2018). To enhance coalescent modelling by representing genetic diversity (Edwards et al. 2016), samples were chosen from distant populations and diverse wing pattern races as available. The total dataset included 145 individuals across 40/45 species of *Heliconius*, 6/12 *Eueides* and the monotypic *Dryadula* and *Agraulis* (S1 Table).

Sequencing was performed using the Illumina technology (S1 Table) with 100 bp paired-end reads, insert sizes of 250 - 500 bp and read coverage from 12x to 110x. In case of 13 new samples, DNA was extracted with the DNeasy Blood and Tissue kit (Qiagen) from 30-50 µg of thorax tissue homogenised in buffer ATL using the TissueLyser (Qiagen); purified by digesting with RnaseA (Qiagen); and quantified on a Qubit v.1 spectrophotometer (LifeTechnologies). The Beijing Genomics Institute constructed and sequenced whole-genome libraries on a HiSeq 2500 with 500 bp insert size to ∼50x coverage (S1 Table).

### Read mapping and genotyping

Raw reads were checked using FastQC v0.11 (Andrews 2014) and mapped to the *H. melpomene melpomene* reference (Heliconius Genome Consortium 2012) with BWA v6(Li & Durbin 2009), followed by refinement with Stampy v1.0.18 (Lunter & Goodson 2011). Aligner parameters were based on empirical tests (Nadeau et al. 2013; Davey 2013) and the age of divergence from the reference (Kozak et al. 2015) (S2 Table). Alignments were sorted with Samtools(Li et al. 2009), de-duplicated with Picard v1.112 (Fennell 2010) and re-aligned in Genome Analysis Toolkit v3.1 (GATK) (McKenna et al. 2010; DePristo et al. 2011). SNPs were called separately across samples at sites with coverage >4x and quality >20 using the GATK UnifiedGenotyper(van der Auwera et al. 2013). The liberal values that increase calling sensitivity were chosen as appropriate for a phylogenetic analysis, where a few individual SNPs are not viewed as evidence in isolation. Species genotypes were merged using Bcftools v1 (Li et al. 2009) and assessed with an in-house Python script (Martin et al. 2013) (evaluateVCF-03.py (Martin 2017)). We identified 126,865,683 individual SNPs, including 5,483,419 in the exome. The autosomal matrix of exonic, biallelic, non-singleton SNPs genotyped in all individuals contained 122,913 variants (Supplementary Methods).

*Commands for genotyping and phylogenetic software are listed in Supplementary Methods.*

### Exome alignments and gene trees

Protein-coding genes can be effectively treated as discrete markers for multilocus phylogenetics(Edwards et al. 2016) with lower level of homoplasy than the non-coding sequence(Brawand et al. 2014). We minimized paralogy by narrowing the gene set to 1:1:1 orthologs between *H. melpomene, Danaus plexippus* (Zhan et al. 2011) and *Bombyx mori* (Xia et al. 2004) identified by OrthoMCL (Li et al. 2003; Heliconius Genome Consortium 2012). 6848 autosomal gene alignments, excluding those found on the colour pattern scaffolds (Heliconius Genome Consortium 2012), and 416 Z-linked single-copy genes were extracted using the tabular “calls file” format [gene_fasta_from_reseq.py (Martin 2017)]. TrimAl v1.2 (Capella-Gutiérrez et al. 2009) was used to remove the taxa for which 50% of residues did not overlap with at least 50% of the other sequences, while uninformative fast-evolving regions were excluded by Block Mapping and Gathering with Entropy (BMGE) (Criscuolo & Gribaldo 2010). Individual ML gene trees were estimated in FastTree v2.1 (Price et al. 2010) with parametric aLRT nodal support (Anisimova & Gascuel 2006).

### Incongruence in the data

To assess the level of topological variation in the gene trees, the average normalised Robinson-Foulds distance (RF) (Robinson & Foulds 1981) between all pairs of phylogenies was computed in PAUP* v4 (Swofford 2002). The calculation was repeated after trimming to 57 individuals representing species or highly distinct subspecies (e.g. the three geographic clades of *H. melpomene*), thus eliminating the noise from lack of intraspecific resolution. The 57 samples were chosen based on the quality of the genotype calls (S1 Table). In addition, the average distance between all possible triples of taxa represented in full gene trees - a measure similar to the RF - was calculated in MP-EST (Liu et al. 2010). To identify incongruent nodes, 50% Majority Rule consensi were calculated from the trimmed gene trees and the relative support for branches was evaluated under the Internode Certainty criteria (IC/ICA/TCA), which compare the support for a branch and for all alternatives found in the distribution (Salichos & Rokas 2013; Salichos et al. 2014). The effect of poor resolution in some gene trees was accounted for by repeating the procedure with the 1000 most resolved and supported phylogenies. To quantify the gene-species tree disagreements, we computed what proportion of gene trees contain the quartets found in the ASTRAL-III phylogeny (–*t1*) (Sayyari & Mirarab 2016).

### Species trees

Naïve “total evidence” phylogenies were estimated from concatenated genome-wide SNPs in RAxML v8 with 100 bootstrap replicates, GTR+Γ model and ascertainment bias correction (Stamatakis 2014). The history of the matriline was approximated from the whole-mitochondrial alignment with partitions determined by PartitionFinder v1.1 (Lanfear et al. 2012). To verify a previous dating of the radiation(Kozak et al. 2015), an ultrametric Minimum Evolution tree was estimated from the autosomal SNPs using relative rate comparison in RelTime (Tamura et al. 2012; Tamura et al. 2013), constraining intervals around the age of the *Heliconius-Eueides* (17.0 - 20.0 MA) and *Heliconius-Agraulis* (25.0 - 28.0 MA) as determined for the Nymphalidae phylogeny (Wahlberg et al. 2009).

Two Multispecies Coalescent (MSC) approaches were used to estimate the species tree under the assumption of Incomplete Lineage Sorting. MP-EST v1.4 maximises a pseudo-likelihood function over the distribution of taxon triples extracted from gene tree topologies(Liu et al. 2010). 145 individual tips were assigned to species *a priori* (S1 Table), treating Western and Eastern clades of *H. melpomene* and *H. erato* as distinct (Nadeau et al. 2013; van Belleghem et al. 2017). Bootstrap support was evaluated by repeating 100 times with random samples of 500 gene trees. Since MP-EST may be misled by errors in gene tree reconstruction(Mirarab & Warnow 2015), we compared the results with ASTRAL-III, a fast quartet method that deals with polytomies and low support values in the input (Zhang et al. 2017).

### Modelling hybridisation

Following the identification of problematic nodes based on ICA and MSC trees, we applied three radically distinct approaches to disentangle ILS from hybridization (Hahn & Nakhleh 2016; Peter 2016; Scornavacca & Galtier 2016). First, we determined the Ancestral Recombination Graph (ARG) in TreeMix v1.13 by identifying the pairs of taxa sharing more than the expected proportion of allelic variation (Pickrell & Pritchard 2012). Relations between major lineages and species were inferred from allele frequency data computed in PLINK! v2 (Purcell 2009; Chang et al. 2015) after Linkage Disequilibrium prunning (Gaunt et al. 2007), removing sites with less than 95% complete data, and grouping SNPs into blocks of 100. Models were fitted with zero, 10 or 20 migration events, treating *H. aoede* as an arbitrary outgroup for rooting.

Second, we tested the fit of the data to the bifurcating tree ideal by calculating the *f4* statistic (Reich et al. 2009), a difference in allele frequencies of two taxa pairs that can be interpreted as a measure of treeness (Peter 2016), similar to the *D* statistic (Durand et al. 2011). The *f4* was calculated for all possible quartets of taxa in each clade using the *fourpop* routine in TreeMix on the SNP data described above, determining the statistical significance of the results by jacknifing.

We modeled both hybridization and incomplete coalescence under the coalescent network framework implemented in Phylonet v3.5 (-*InferNetwork_MPL*). Networks were computed under the Maximum Pseudo-Likelihood (MPL) criterion from the 6724 *Agraulis*-rooted autosomal phylogenies, considering only nodes with support >0.8 (*-b 0.8*) and starting with the MP-EST species. The analysis was conducted separately for each clade among *Eueides, H. melpomene, H. erato, H. sara, H. hecuba* and *H. egeria*. To detect ancient admixtures between basal branches of the tree, we conducted an analysis where all individuals of each clade were treated as a single “species”. The optimal network was determined by calculating the BIC from the Maximum Likelihood and the number of lineages, admixture edges and gene trees in each model.

*Heliconius hecalesia* and *H. clysonymus* were difficult to place with confidence in the phylogenies, and clustered with *H. hortense* at the *B/D* locus. To test for gene flow between these species and relatives we calculated the *D* statistic (Durand et al. 2011), determining significance by block jacknifing (Martin et al. 2015). Specific tests were conducted for gene flow between *H. hecalesia* (recipient, P2) and *H. clysonymus, H. hortense, H. telesiphe* or species from the *H. sara* clade (donor, P3). The sister species P1 was either the allopatric *H. erato* from French Guiana, or the parapatric *H. erato* from Amazonia, and the outgroup was always *H. melpomene*. To test the possibility that the lack of resolution at one node in the larger phylogeny (Fig. 1; S1 Figure) is caused by ancient admixture prior to the formation of current species, we conducted tests treating entire clades as taxa (Table 3). The extent of similarity between species clusters was illustrated with PCAs of genome-wide variation, calculated for the *H. erato/sara* clade in the R package *adegenet*(Jombart & Ahmed 2011). A chronogram for the putatively introgressing *B/D* red pattern intervals (HE670865: 320-380 and 410-440 kbp) was estimated with RelTime.

### Introgression at the colour pattern loci

Out of 4309 scaffolds, the five fragments containing introgressing loci associated with aposematic wing phenotypes (Table 2) were treated separately. Each scaffold alignment was partitioned into windows of 20 kbp, sliding by 10 kbp, discarding windows with less than 1000 bp of data [script sliPhy3.py(Martin 2017)]. The topology for every window was reconstructed with FastTree, tested for significant differences from the MP-EST species tree using the SH test (Shimodaira & Hasegawa 1989) in RAxML and inspected visually. In order to understand precisely how the *Hmel1* reference corresponds to the specific red control loci of *H. erato* (Supple et al. 2013), the *B/D* scaffold HE670865 was aligned against the *H. erato B/D* BACs (Papa et al. 2008) with mLAGAN (Brudno et al. 2003).

#### ABBREVIATIONS

ARG: Ancestral Recombination Graph
BIC: Bayesian Information Criterion
IC: Internode Certainty
ILS: Incomplete Lineage Sorting
LD: Linkage Disequilibrium
ML: maximum likelihood
MSC: Multispecies Coalescent
SNP: single nucleotide polymorphism

## ACKNOWLEDGMENTS

We thank the governments of Peru, Ecuador and Suriname for permits to collect butterflies. Kanchon Dasmahapatra, James Mallet and Camilo Salazar provided advance access to genomic data and helpful comments. Analyses were conducted on a machine made available by Aylwyn Scally, clusters at the School of Life Sciences operated with help from Jenny Barna, and the STRI *Plato* server run by Eugenio Valdes. Richard Nichols and John Welch examined an early draft of the manuscript.

## MAIN TABLES AND FIGURES

## SUPPLEMENTARY METHODS

### Software command lines and parameters

#### Mapping

Settings for the most divergent genomes were based on experiments with *Eueides lybia* and *Acraea encedon* libraries (BWA mismatch numbers *k*={2, 3}; *l* ={15-35}; Stampy substitution rate ={0.1-0.2}). The proportion of properly paired reads did not change by more than 4% among the combinations, and most conservative values were used as listed in S2 Table.

#### GATK UnifiedGenotyper calling

GenomeAnalysisTK.jar -T UnifiedGenotyper -R Hmel1.1.fasta -I individual.bam -o individual_variants.vcf --output_mode EMIT_ALL_SITES --downsample_to_coverage=250 --baq=CALCULATE_AS_NECESSARY

#### TrimAl v1.2 alignment trimming for of sites with most data missing

trimal -in alignment.fasta -out alignment.trimal.fasta -fasta -resoverlap 0.5 -seqoverlap 50

#### Removal of uninformative sites by Block Mapping and Gathering with Enthropy (BMGE)

BMGE.jar -i alignment.trimal.fasta -t DNA -m DNAPAM100:1 -ory alignment.RY.phy -s NO -g 0.5

#### FastTree v2 ML gene tree estimation

FastTree -nt -gtr -pseudo -spr 4 -gamma -log alignment.ft.log $f > alignment.ft.tre

#### RAxML v8 ML supermatrix tree estimation

raxmlHPC-PTHREADS-SSE3 -T 10 -f d -p 123 -m GTRGAMMA -s supermatrix.fasta -n supermatrix.ML.tre

#### RAXML v8 Internode Certainty calculation

raxmlHPC-PTHREADS-AVX –T 10 -f i -m GTRCAT –t supermatrix.tre -z gene.trees -n ica.analysis

#### MP-EST v1.4 control file lines

gene.trees

1

-1

6848 52

{mapping of individuals to species}

0

*Astral-III command:*

astral.5.5.9.jar -i in.tre

astral.5.5.9.jar –i gene.trees –t 1 –a mapping.to.species.clust –o astral.tre

#### Plink v2 SNP filtering

plink --file treemix.plink –ld –freq –noweb –missing 0.05 --within mapping.to.species.clust

#### TreeMix v1.13 ARG with one admixture

treemix -i treemix.calls.gz –m 1 –k 100 -root H_erato -o H_melpomene_clade

#### TreeMix v1.13 Reich’s *f*_*4*_ statistics

fourpop -i treemix.calls.gz –m 1 –k 100 > f4.out

*Phylonet Maximum Pseudo-Likelihood network with one hybridisation event:* InferNetwork_MPL (Autosomal_tree_1-Autosomal_tree_6725) 1 -n 3 -pl 12 -x 10 -s mpest_start_tree -di -b 0.8

### Results of read mapping across the genus to the *Hmel v1* reference

New Whole Genome Illumina resequencing data were produced successfully for 11 species, extending the sampling of the *H. sara* clade and *Eueides*. For all Illumina samples, the mean per-base quality was high along the entire reads (Q>28) and all the quality filters were passed. The number of reads ranged from 94,857,214 to 194,528,700 per individual, corresponding to an expected coverage of 34x to 71x (S3 Table).

Most clades within *Heliconius* (as defined in Brown 1981 and Kozak et al. 2015) were sampled thoroughly, except the inclusion of only one out of four species in the subgenus *Neruda* (see Chapter 2 for taxonomy). Among the 23 *Heliconius* species with multiple samples, the number of individuals ranged from two to 25, the number of geographically distinct populations was between one and 12, and the number of colour pattern races ranged from one to 14 (S1 Table).

The depth and number of mapped reads decreased predictably with increasing divergence from the reference (Table 5.3). In case of the MCS clade, more than 90% of reads mapped to the *Hmel1*, and most were properly paired with their mate. The number of mapped reads fell steeply for more distantly related taxa, averaging 58.6% for *H. erato,* 43.2% for *Eueides* and around 30% for the outgroups. As the protein coding sequence comprises 5% of the *H. melpomene* genome (Heliconius Genome Consortium 2012), the data obtained for these more distantly related species is mostly non-coding sequence. The non-heliconian outgroup *Acraea encedon* (Acraeini) and the *H. doris* sample 8684 were excluded due to low effective coverage (respectively 1.5x and 3.4x) manifested in poor quality of inspected alignments. Regardless of divergence from the reference the mapping was the highest for samples with expected coverage >40x and usually low at <15x.

The number of SNPs follows similar trends, The highest number of credible variants was called in the MCS clade (S3 Table). More distant taxa showed naturally higher variation from the reference, which simultaneously reduced the overall number of mapped sites. In total, 126,865,683 SNPs were identified, a number driven primarily by taxon-specific differences from the reference, but also by high variability in the densely sampled MCS group. For instance, private SNPs constituted 30% of the total and 11.5*10^6^ private variants were discovered just in the *H. melpomene* samples. 5,483,419 SNPs were found in the exome. Overall, these findings are consistent with previous reports for the MCS group (Martin et al. 2013) and the monarch butterflies *Danaus* (Zhan et al. 2014).

The transition/transversion ratio for the MCS clade is a low 1.25, but similarly small values were previously found in the noncoding sequences of the cricket *Podisma pedestris* (Keller et al. 2007). The bias increases with divergence, reaching 1.43 for *Eueides* and 1.50 for *Dryadula*, most likely due to a higher proportion of CDS in the recovered total (Table 5.3). Such variation in the Ts/Tv bias is observed in the human data, where the Ts/Tv equals 2.1, but increases to 3.0 in the exome (1000 Genomes Initiative 2015).

## SUPPLEMENTARY TABLES

**S1 Table. Specimens – XLS file.**

**S2 Table.** Empirically adjusted parameters for the short read alignment to the *H. melpomene* reference. *k*=maximum number of mismatches per 100 bp; *l*=minimum stretch of identical sequence necessary to map; *subRate*=expected nucleotide divergence.

| Clade | Divergence (Myr) | BWA *k* | BWA *l* | Stampy *subRate* |
| --- | --- | --- | --- | --- |
| *H. melpomene* | <1.5 | 2 | 32 | 0.03 |
| *H. cydno* | 2.0 | 2 | 32 | 0.04 |
| Silvaniforms | 4.0 | 2 | 32 | 0.05 |
| *H. erato* | 12.0 | 2 | 25 | 0.1 |
| *Eueides, Agraulis, Acraea* | >18.5 | 2 | 25 | 0.1 |

**S3 Table.** Mapping quality and number of SNPs decrease with divergence from the reference. Statistics for the BWA/Stampy read mapping of Illumina 100 bp paired-end reads to the *H. melpomene* reference, averaged by clade *sensu* Brown 1981. Values were calculated for sites with quality score 20 or higher. Ranges reported in parentheses. *Percentages of reads mapped reported for the best sample.

| Clade | Species | Samples | Coverage x | Reads mapped* | Reads properly paired* | # SNPs | Biallelic SNPs | Singletons | Ts/Tv |
| --- | --- | --- | --- | --- | --- | --- | --- | --- | --- |
| *H. melpomene* | 1 | 25 | 27.33 (5.19-85.58) | 316,265,364 (94.55%) | 262,954,142 (78.62%) | 32,788,205 | 30,615,977 | 11,497,421 | 1.27 |
| *H. melpomene*/ *H. cydno* | 5 | 48 | 26.01 (5.19-100.01) | 388,402,453 (93.80%) | 309,082,266 (74.64%) | 48,793,385 | 44,279,711 | 16,910,938 | 1.26 |
| Silvaniform (*H. numata*+relatives) | 8 | 28 | 21.82 (8.05-33.69) | 142,606,506 (90.21%) | 97201540 (61.49%) | 57,561,620 | 50,888,801 | 21,259,708 | 1.24 |
| *H. wallacei* | 3 | 4 | 10.16 (5.32-16.80) | 81,888,086 (67.03%) | 31084824 (25.45%) | 17,425,503 | 16,832,134 | 3,341,464 | 1.28 |
| *H. doris* | 4 | 6 | 8.46 (3.43-22.85) | 172,414,557 (72.19%) | 70,011,956 (29.31%) | 35,334,399 | 32,579,459 | 5,613,625 | 1.25 |
| *H. aoede* | 1 | 1 | 14.36 | 79,473,860 (61.97%) | 31,150,826 (24.29%) | 8,581,604 | 7,257,197 | 8,581,604 | 1.27 |
| *H. erato* | 7 | 33 | 10.57 (5.11-16.55) | 138,223,134 (64.01%) | 49,770,828 (23.05%) | 33,988,575 | 30,643,397 | 7,578,237 | 1.32 |
| *H. sara* | 12 | 17 | 10.02 (5.55-19.07) | 138064019 (58.63%) | 48,057,198 (20.41%) | 26,791,561 | 24,831,585 | 4,599,643 | 1.30 |
| *Eueides* | 6 | 6 | 6.07 (4.90-7.54) | 72,571,204 (43.22%) | 15,904,356 (9.47%) | 12,905,223 | 12,005,724 | 1,966,268 | 1.43 |
| *Agraulis* | 1 | 1 | 7.26 | 58,461,652 (30.57%) | 14,577,290 (7.62%) | 4,949,437 | 4,459,801 | 4,949,437 | 1.47 |
| *Dryadula* | 1 | 1 | 3.61 | 27,336,635 (30.85%) | 6,130,764 (6.92%) | 4,141,846 | 3,975,796 | 4,141,846 | 1.50 |
| **TOTAL** | **48** | **145** | **17.4 (3.43-100.01)** | **n/a** | **n/a** | **126,865,683** | **90,646,525** | **38,070,723** | **1.29** |

**S4 Table. TreeMix and Phylonet Model selection –XLS file.**

**S5 Table.** Basic statistics for the autosomal and Z-linked protein-coding gene alignments. Relatively short sequences were removed with TrimAl and uninformative sites were deleted by Block Mapping and Gathering with Entropy (BMGE). Range of lengths after trimming in parentheses.

| Parameter | Autosomal | Z-linked |
| --- | --- | --- |
| # alignments | 6848 | 416 |
| # *Agraulis*-rooted alignments | 6724 | 406 |
| Taxa after TrimAl | 144.53 | 144.75 |
| Length before BMGE (bp) | 1399 | 1633 |
| Length after BMGE (bp) | 1387 (60-15,921) | 1627 (210-11,979) |
| Missing data | 4.0 % | 3.6 % |
| Ambiguous sites | 1.3 % | 0.3 % |
| GC content | 42.9 % | 44.6 % |
| Interspecific pairwise identity | 90.8 % | 89.7 % |
| Gene tree length | 0.7128 | 0.7416 |

## SUPPLEMENTARY FIGURES

**Figure S1.**
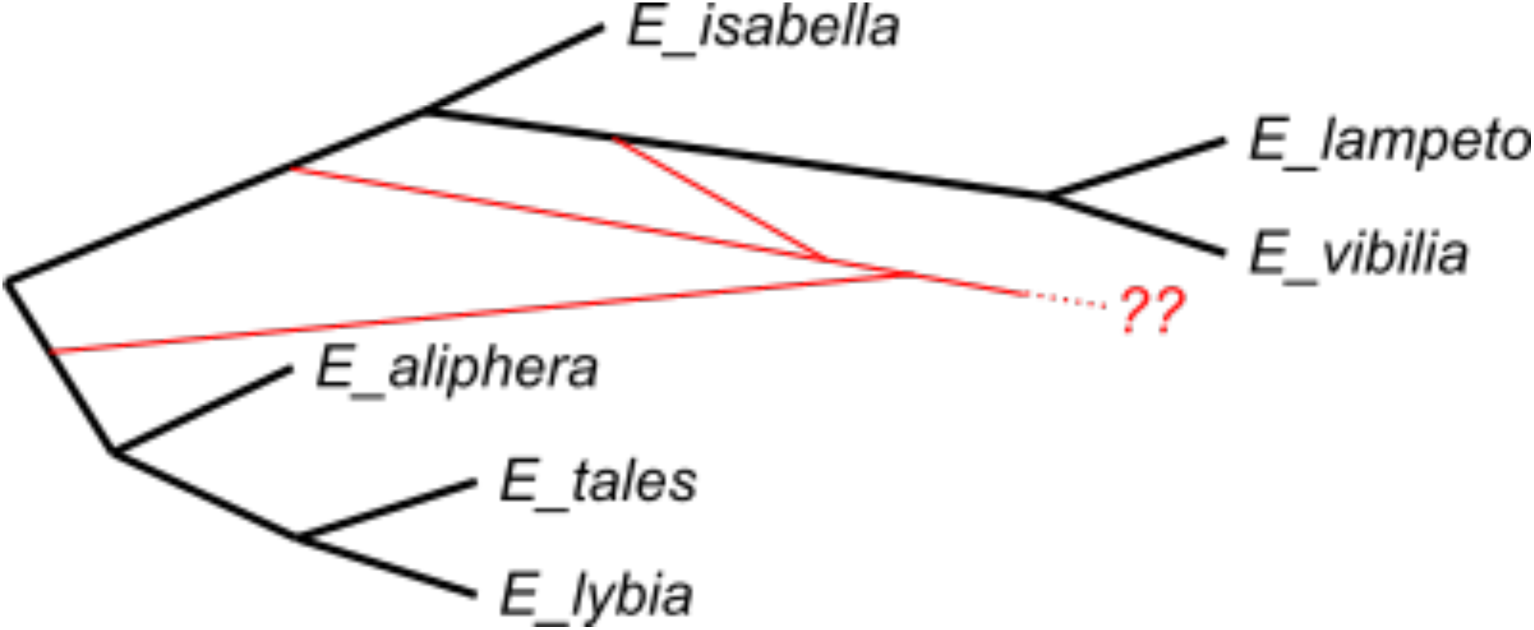
Inconsistent estimates of gene flow in a representation of *Eueides* (6/12 species), the sister group of *Heliconius*. MPL network from the autosomal gene trees. The vertices of multiple branches may represent an artefactual “ghost lineage” corresponding to missing species.

**Figure S2.**
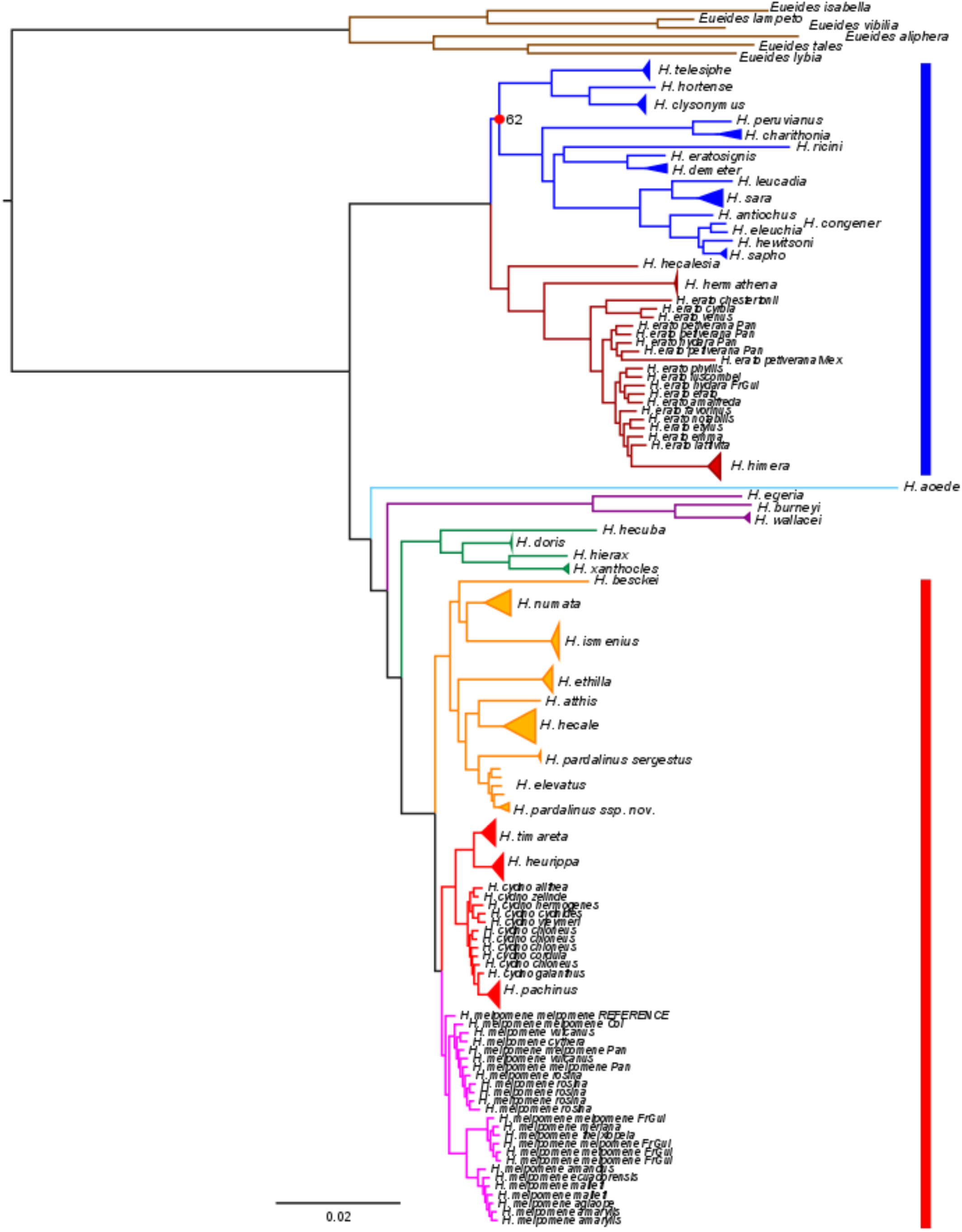
High support for a concatenation tree based on autosomal SNPs. All nodes in the Maximum Likelihood (RAxML) phylogeny have a bootstrap support of 100, except for the split labeled with a red dot (62/100). Most intraspecific samples collapsed. Branches coloured by clade as defined in Chapter 2: brown – *Eueides*; red – *H. sapho* clade; navy – *H. erato* clade; blue – *H. aoede* clade (formerly *Neruda*); green *H. doris* clade; violet – *H. wallacei* clade; orange – Silvaniforms; red – *H. cydno* and cognates; pink – *H. melpomene*.

**Figure S3.**
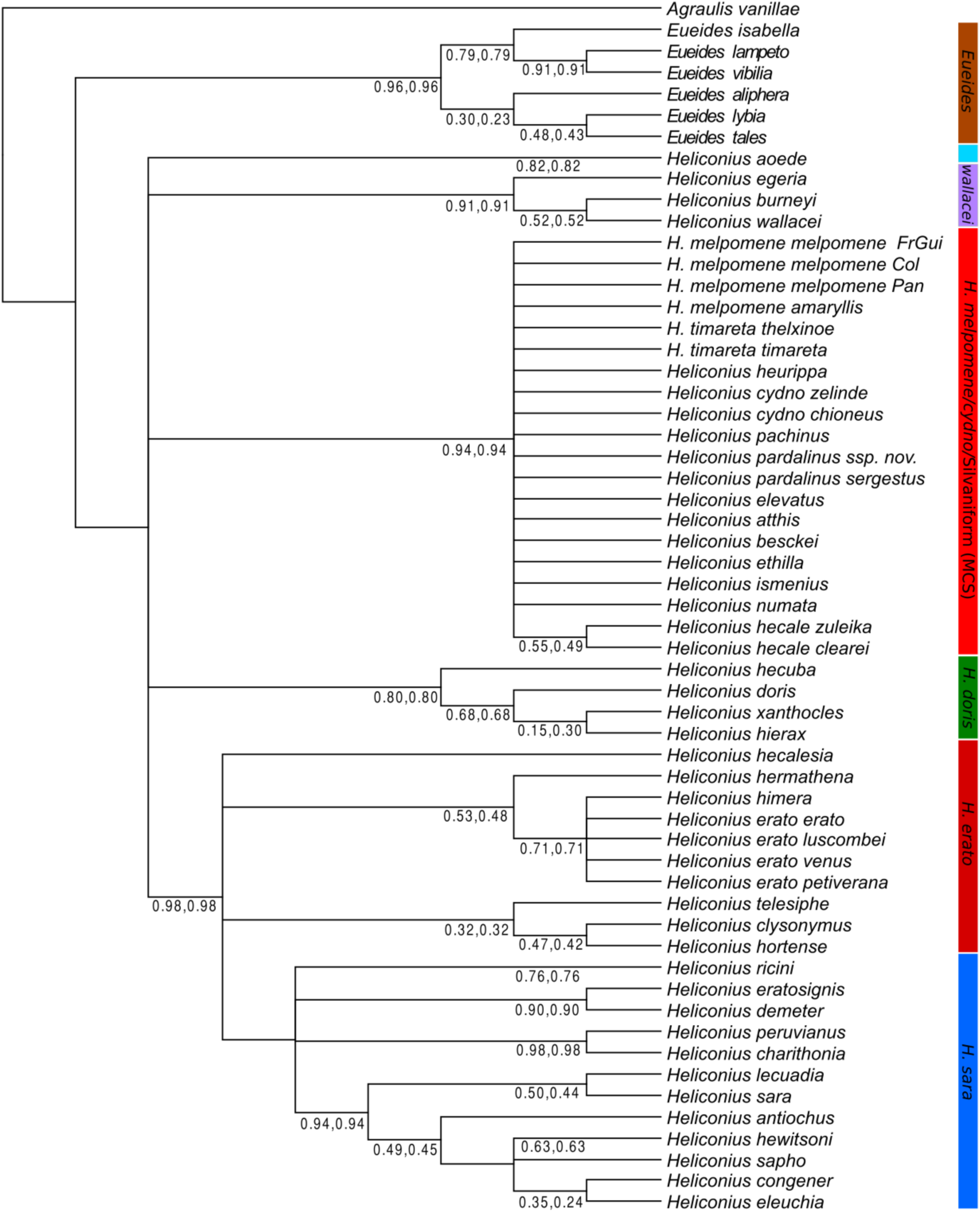
Autosomal gene trees disagree at most nodes. 50% Majority Rule Consensus tree based on 6724 Agraulis-rooted gene trees. Branch labels indicate the IC and ICA support (Salichos et al. 2014). An IC=0.0 means that a given node has an equally frequent alternative in the distribution of gene trees, whereas IC=1.0 means that all trees contains this node. FrGui: French Guiana; Col: Colombia; Pan: Panama.

**Figure S4.**
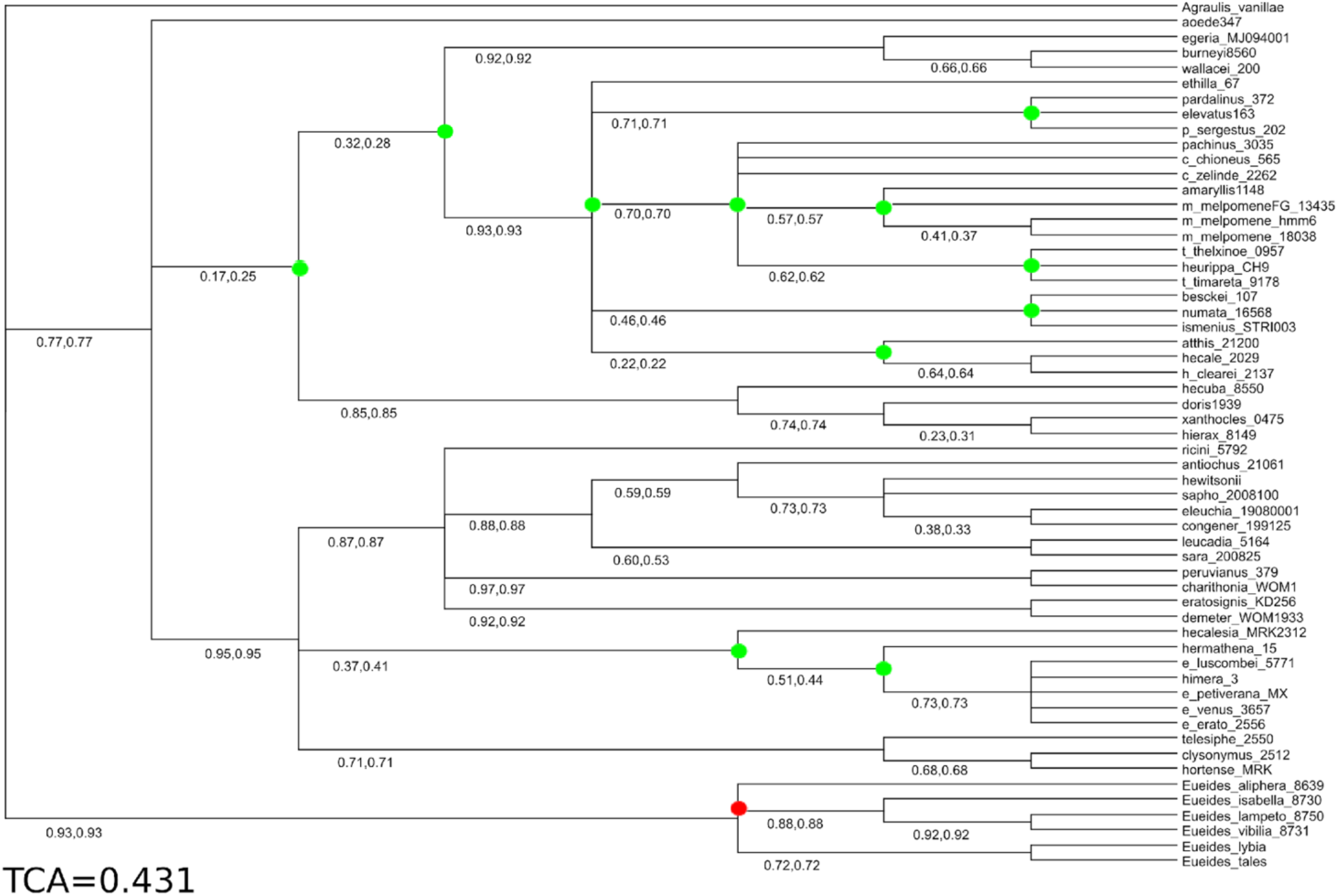
More interspecific genetic isolation at the Z chromosome than at the autosomes. 50% Majority Rule Consensus the Z-linked gene trees with IC/ICA support values indicated. Green dots indicate nodes unresolved in the autosomal 50% MRC, red dots nodes conflicting with the autosomal tree.

**Figure S5.**
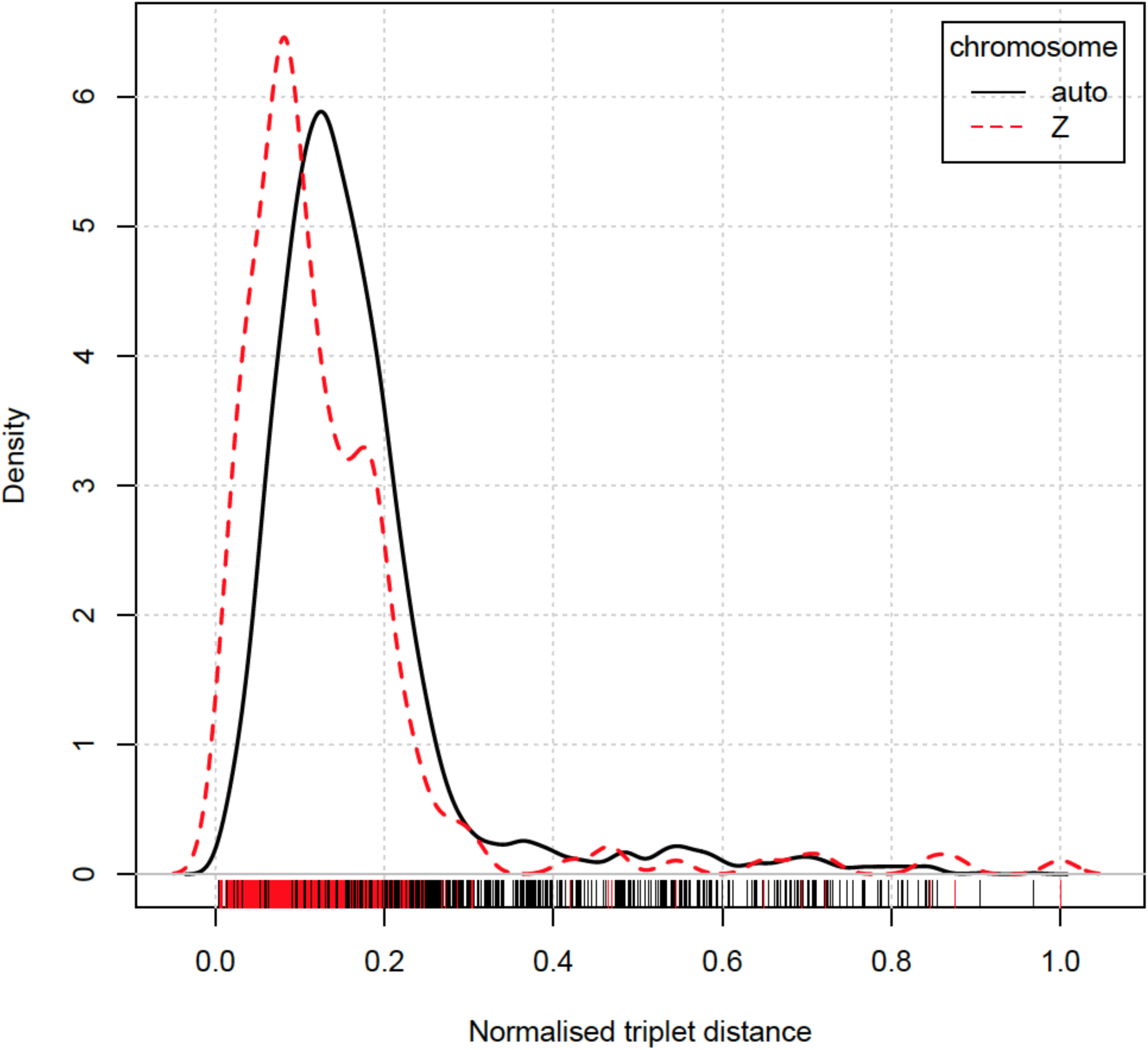
Autosomal and Z-linked genes are almost equally conflicted with the species tree. A smoothed distribution of the normalised triplet distance between the gene trees and the MP-EST species tree.

**Figure S6.**
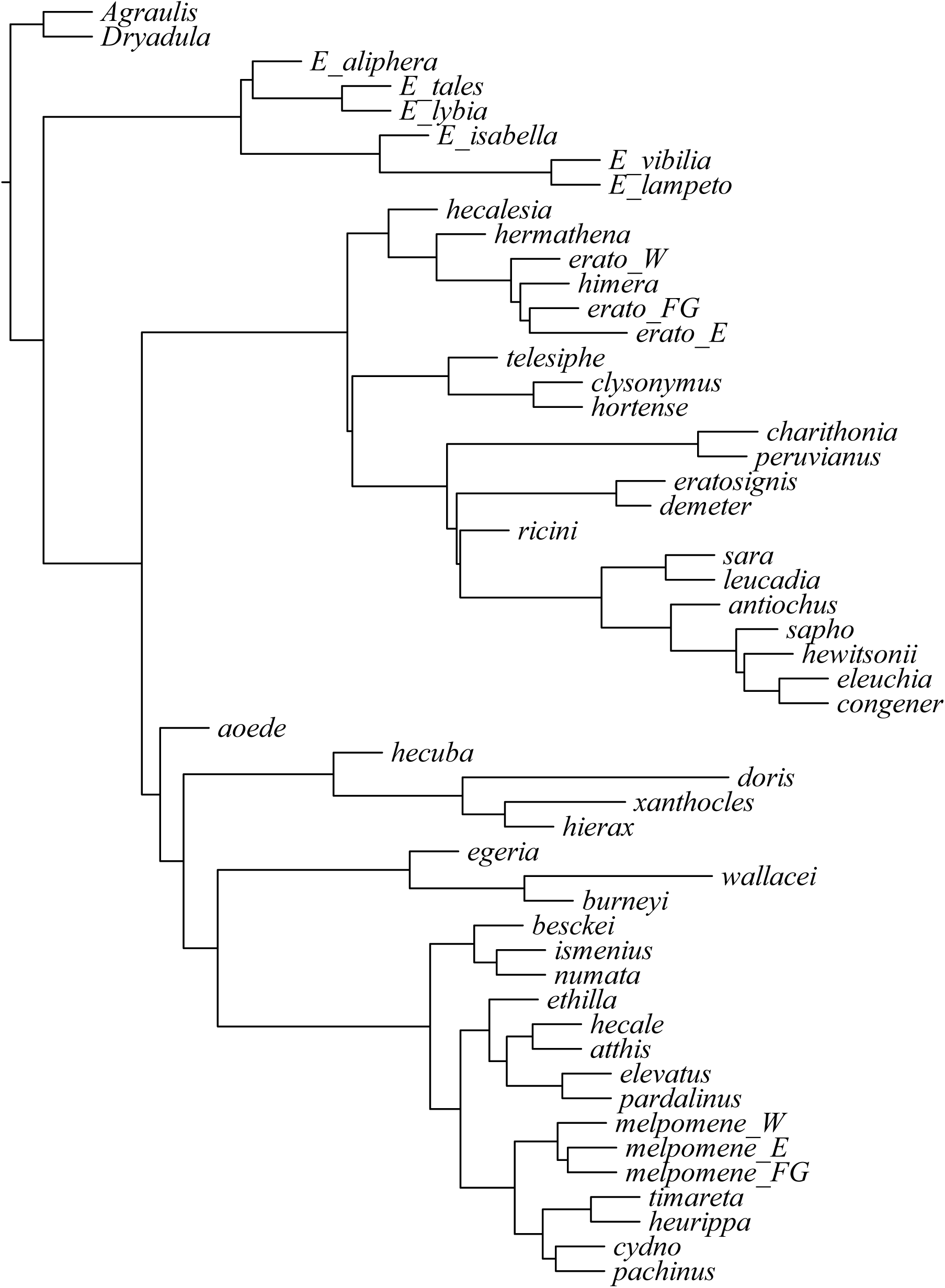
Resolved and supported ASTRAL-III phylogeny for the Z (sex-linked) chromosome.

**Figure S7.**
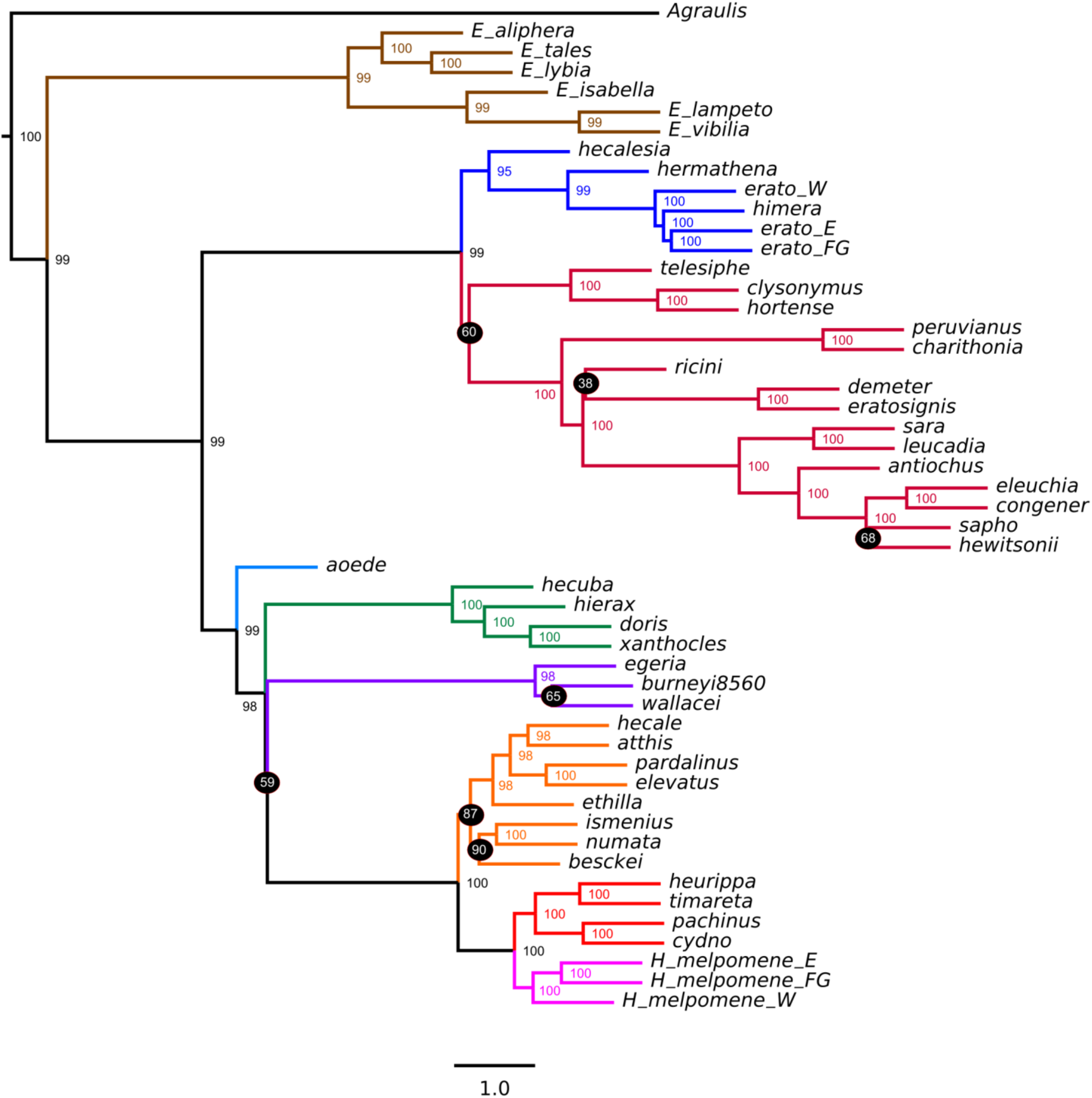
Incomplete lineage sorting at the autosomal loci. A multispecies coalescent tree estimated from the 6848 autosomal CDS gene trees under the MPEST pseudolikelihood model shows lack of resolution at several nodes. Branch lengths in coalescent units, terminal branch lengths arbitrarily set to 1.0. Bootstrap support values indicated.

**Figure S8.**
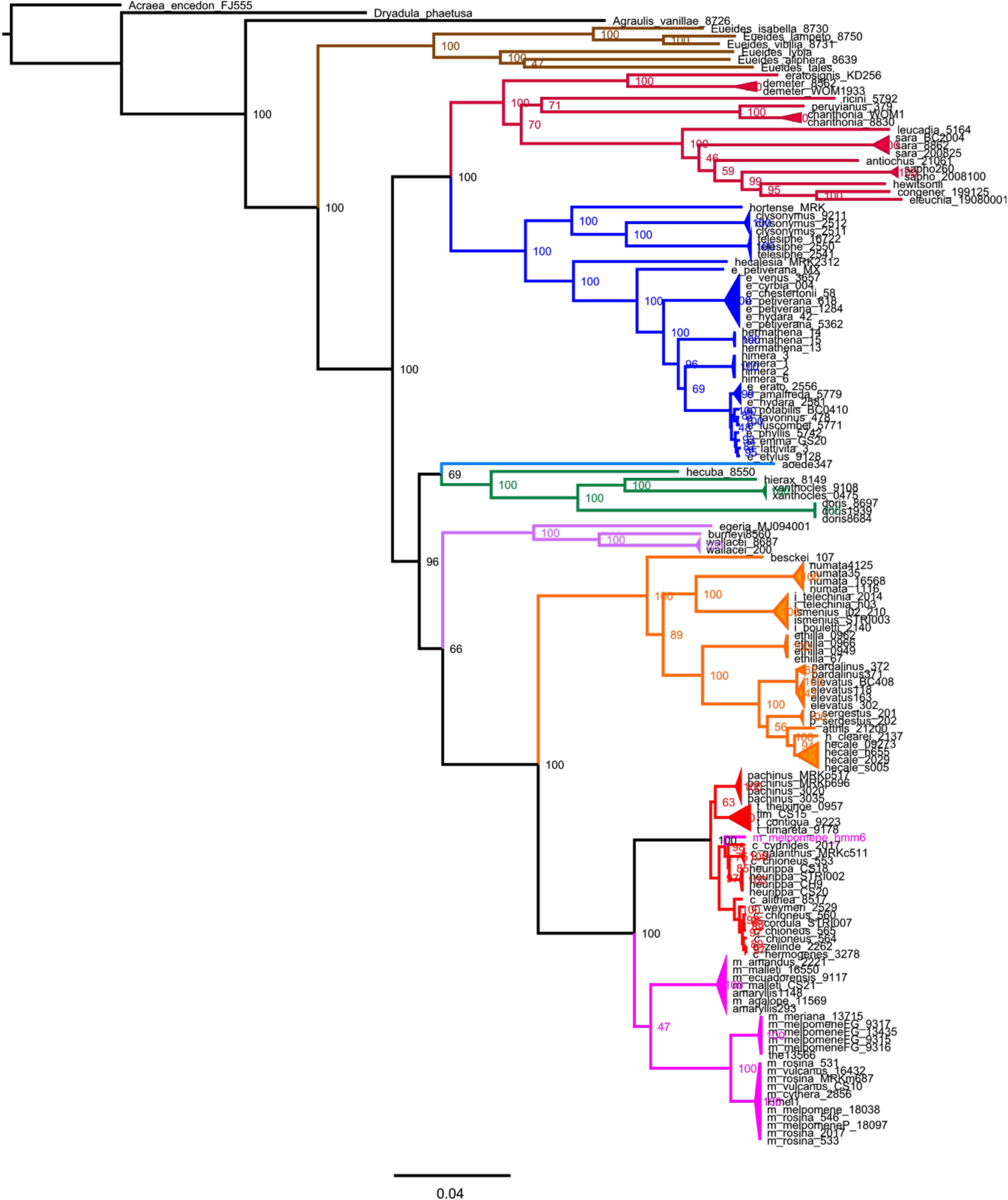
Whole-mitochondrial sequences “resolve” the radiation of *Heliconius*, although they conflict with autosomal and sex-linked signals. A Maximum Likelihood tree (RAxML) with bootstrap support values. Colours indicate major clades *sensu* Brown (1981). Scale bar in substitutions per site.

**Figure S9.**
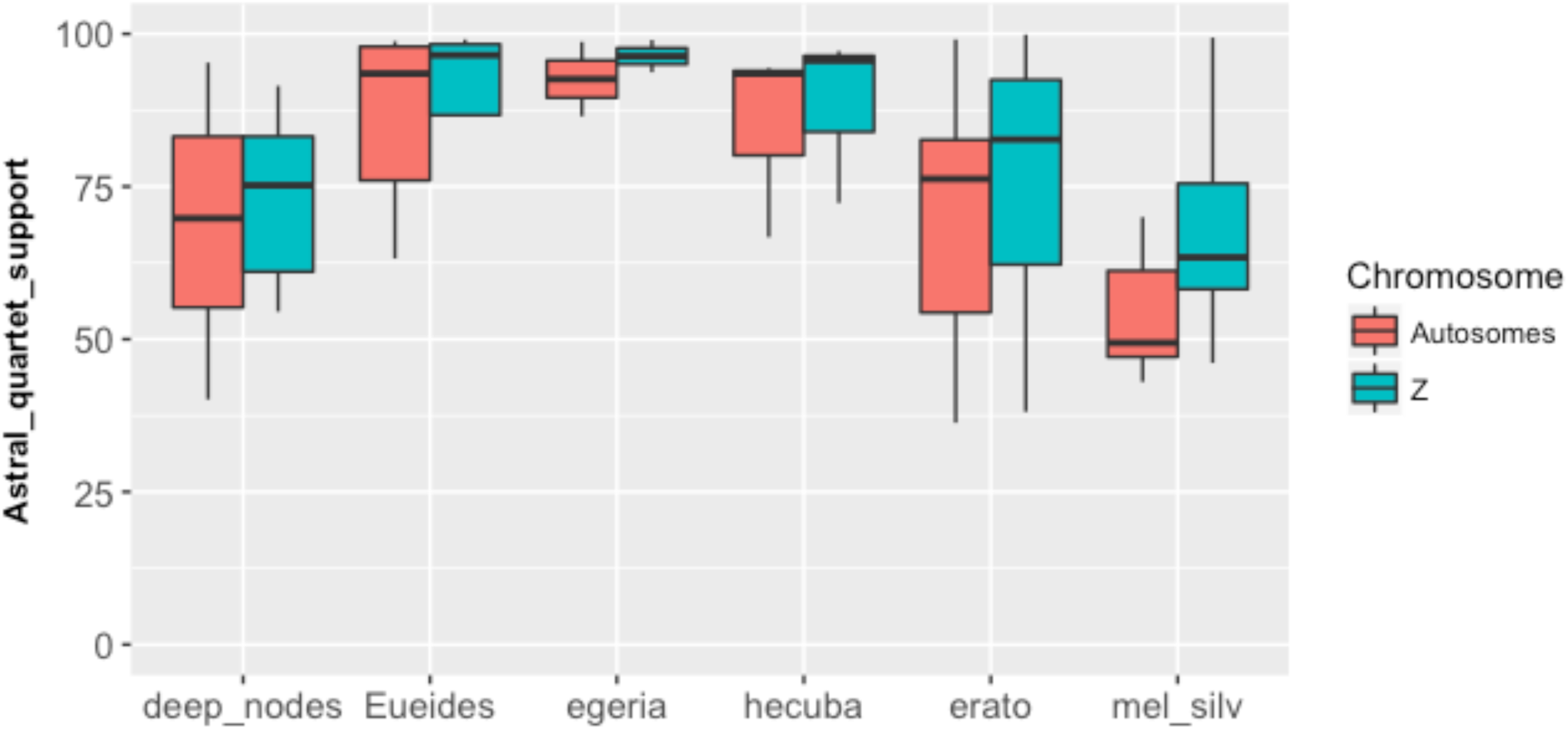
Gene tree incongruence is the highest in the *H. melpomene/*silvaniform clade. Incongruence is higher at the autosomes than the Z sex chromosome. Mean proportion of gene tree quartets supporting all nodes as calculated in ASTRAL is plotted for all groups of *Heliconius*; *Eueides*; and the deep nodes that link them. Higher support values indicate less incongruence between individual gene trees.

